# In situ structural analysis reveals membrane shape transitions during autophagosome formation

**DOI:** 10.1101/2022.05.02.490291

**Authors:** Anna Bieber, Cristina Capitanio, Philipp S. Erdmann, Fabian Fiedler, Florian Beck, Chia-Wei Lee, Delong Li, Gerhard Hummer, Brenda A. Schulman, Wolfgang Baumeister, Florian Wilfling

## Abstract

Autophagosomes are unique organelles which form *de novo* as double-membrane vesicles engulfing cytosolic material for destruction. Their biogenesis involves a series of membrane transformations with distinctly shaped intermediates whose ultrastructure is poorly understood. Here, we combine cell biology, correlative cryo-electron tomography (ET) and novel data analysis to reveal the step-by-step structural progression of autophagosome biogenesis at high resolution directly within yeast cells. By mapping individual structures onto a timeline based on geometric features, we uncover dynamic changes in membrane shape and curvature. Moreover, we reveal the organelle interactome of growing autophagosomes, highlighting a polar organization of contact sites between the phagophore and organelles such as the vacuole and the ER. Collectively, these findings have important implications for the contribution of different membrane sources during autophagy and for the forces shaping and driving phagophores towards closure without a templating cargo.

## Introduction

Macroautophagy (autophagy hereafter) is a key pathway to maintain cellular homeostasis. In this process, a *de novo* synthesised double membrane vesicle, the autophagosome, engulfs cellular material in response to stress conditions^1^. This culminates in autophagosome fusion with lysosomes (or the vacuole in yeast) to remove and recycle its cargo. Fluorescence microscopy has identified the hierarchical order of the autophagy machinery during autophagosome biogenesis^2,3^. In addition, many of the membrane intermediates have been visualized at low resolution with conventional electron microscopy^4^. These and other methods have revealed that autophagy proceeds in several steps: (1) membrane nucleation, (2) growth of the cup-shaped phagophore, (3) closure and (4) fusion of the autophagosome with the lytic compartment^5^. Meanwhile, pioneering genetic and biochemical studies have revealed key regulators of autophagosome biogenesis^56^. In yeast, nitrogen starvation triggers the first step of phagophore nucleation through assembly of the molecular machinery in the pre-autophagosomal structure (PAS) next to the vacuole^7^. The phagophore is initially formed by fusion of few vesicles carrying the transmembrane protein Atg9^8–10^. It then grows both by fusion of vesicles (e.g. Atg9 or COPII vesicles^11^) and by lipid transfer from the ER through protein complexes such as Atg2/Atg18^12^. Membrane expansion is further driven by conjugation of the ubiquitin-like protein Atg8 to phosphatidylethanolamine (PE) in the phagophore membrane^13^. During growth, the initial membrane disk assumes a characteristic cup shape, a transition that is likely driven by the highly curved and therefore energetically unfavourable phagophore rim^14^. After closure and maturation, the resulting autophagosome fuses with the vacuole, releasing the inner vesicle – now called “autophagic body” – for degradation.

Despite the importance of autophagy and the efforts in deciphering the molecular machinery underlying it^5^, it is still unknown how membranes are organized and transformed on an ultrastructural level during autophagosome biogenesis. *In situ* cryo-electron tomography (cryo-ET) can reveal membrane structures directly in their native cellular environment^15,16^. Yet monitoring the formation of an organelle poses the challenge to capture a rare event with many intermediates along the process. To overcome these hurdles, we combined several strategies to dissect the formation of autophagosomes using cryo-ET: (1) stimulating their formation to increase the abundance of all species involved, (2) using mutants that accumulate intermediates that are naturally short-lived, and (3) fluorescently labelling the autophagy machinery or its cargo to specifically target those structures during focused ion beam (FIB) milling and tomogram acquisition.

Using this approach, we captured the major membrane structures in bulk autophagy for the first time within their native context and at high resolution. Our detailed data analysis provides new insights into the biophysics of autophagosome biogenesis. While we focus here on yeast autophagy, our study highlights the potential of correlative cryo-ET in analysing short-lived cellular structures and provides a general template for studying the formation of organelles.

## Results

### Correlative cryo-ET resolves different steps of autophagosome biogenesis

We used nitrogen starvation to robustly induce autophagy in S*. cerevisiae* cells^17,18^ and employed two alternative fluorescence labelling strategies: (1) overexpressing tagged Atg8 to mark autophagic structures, since Atg8 is conjugated to the phagophore membrane and is present in most stages of autophagy^13^ (Fig. 1a); (2) overexpressing the cargo protein eGFP-Ede1, either alone or in combination with mCherry-Atg8 (Extended Data Fig. 1a-e). Ede1 overexpression leads to accumulation of endocytic machinery proteins at the plasma membrane in a compartment called END (Ede1-dependent endocytic protein deposits), a selective autophagy cargo^19^. Thus, eGFP-Ede1 marks autophagic structures independent of Atg8 overexpression. The starved cells were subjected to a correlative focused ion beam (FIB)-milling and cryo-ET workflow^20,21^: after plunge-freezing, cryo-fluorescence stacks were recorded. Fiducial-based 3D correlation with FIB/SEM and TEM images targeted structures of interest during lamella milling and tilt-series acquisition (Extended Data Fig. 1a-e). The correlation thus identifies and provides evidence for the autophagic nature of the structures in the tomograms (Fig. 1b-d).

**Fig. 1:**
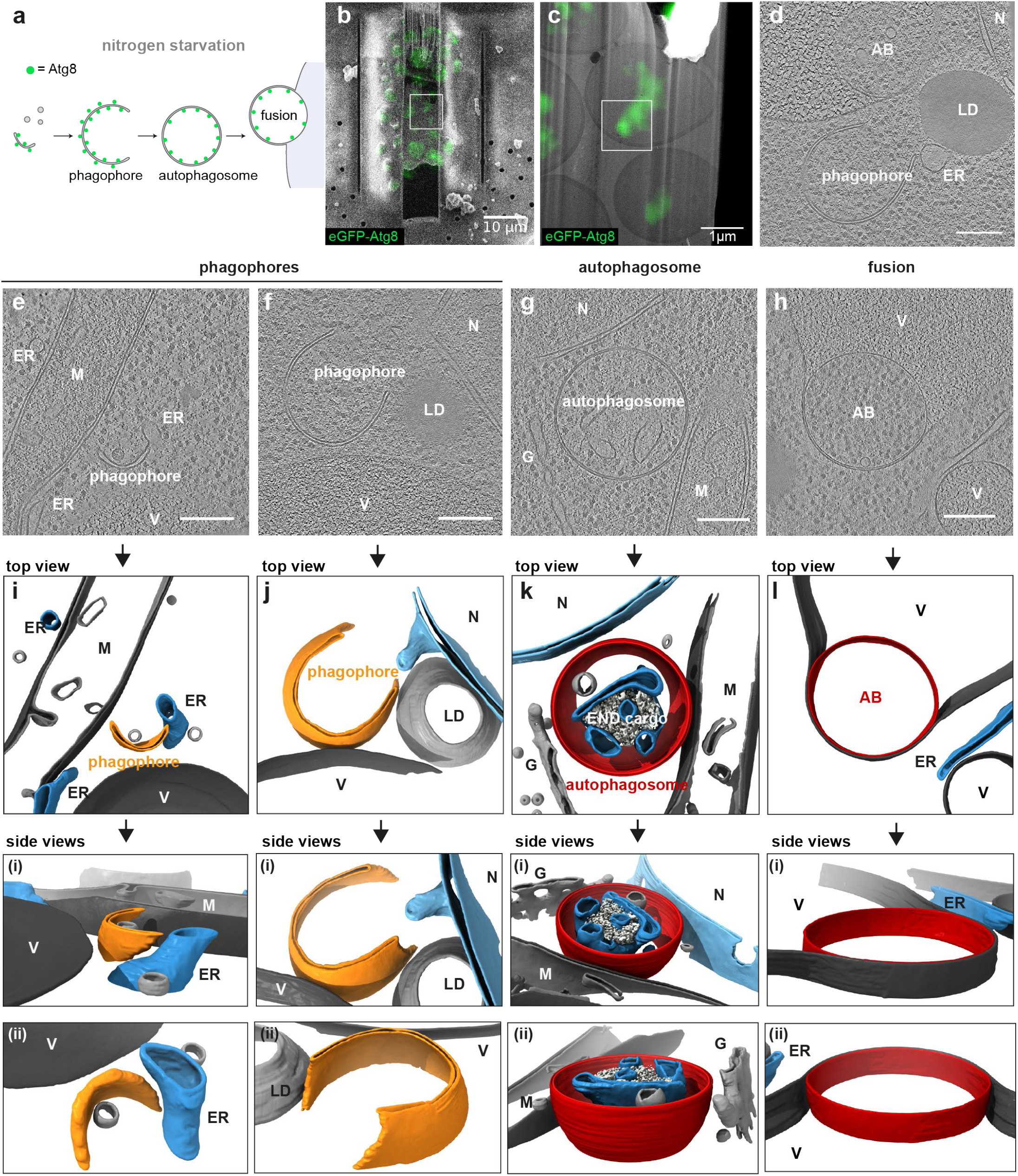
Correlative cryo-ET captures key steps of autophagy in yeast. **a**, Scheme of key autophagy steps and their targeting with eGPF-Atg8. **b** and **c**, SEM and TEM overviews of a lamella, overlaid with correlated eGFP-Atg8 cryo-fluorescence signal (green) used to target the autophagic structures (white boxes). d, Slice through a tomogram acquired in the boxed area in (c) reveals an autophagic body (AB, top) and an open phagophore (bottom). **e-h**, Exemplary tomogram slices and segmentations of the key autophagy steps captured with correlative cryo-ET. All tomogram scalebars: 200 nm. **i-l**, 3D renderings in top and zoomed-in side views (i-ii) of autophagic structures and organelles in the tomograms shown in (**e-h**).

The captured key steps of autophagosome biogenesis (Fig. 1e-h) include early phagophores, in which the double membrane disc is slightly bent to form a small concave structure (Fig. 1e, i and Extended Data Fig. 1a). Next, we frequently observed expanded, cup-shaped phagophores with a clearly visible opening to the cytoplasm (Fig. 1f, j and Extended Data Fig. 1b). Furthermore, we found closed autophagosomes, for which no opening or rim is visible (Fig. 1g, k and Extended Data Fig. 1c), and autophagic bodies, often still partially wrapped by the outer autophagosome membrane fused with the vacuole (Fig. 1h, l and Extended Data Fig. 1d). Importantly, the autophagic structures correlated to the cargo eGFP-Ede1 are indistinguishable from the structures found in cells expressing only eGFP-Atg8 (Extended Data Fig. 1f-h). In total, we collected 35 tomograms of open phagophores, as well as 17 structures without any visible opening. To capture more closed autophagosomes, we created an eGFP-Ede1 mutant strain lacking the Rab7-like GTPase Ypt7, resulting in accumulation of autophagosomes in the cytosol^22^. This strain yielded an additional 25 closed autophagosomes (and 2 phagophores) that closely resembled the ones obtained from the wild type strain (Extended Data Fig. 1f, g, i).

### Cargo templating is not essential for autophagosome formation under bulk conditions

During nutrient starvation, autophagic structures mainly engulf cytosolic ribosomes as shown by earlier EM studies^23^, but can still retain selectivity for specific cargo^24–26^. In line with this, 98 out of 104 autophagic structures contained ribosomes alone or next to selective cargo like the END or the cytoplasm to vacuole targeting (Cvt)^27^ cargo prApe1^28^ (Fig. 2a-c and Extended Data Fig. 2a). Only in few cases, we observed the exclusive uptake of selective cargo (Extended Data Fig. 2b, c). To test if cytosolic cargo clusters to guide phagophore growth during nitrogen starvation, we extracted ribosome positions in the tomograms by template matching and subtomogram averaging (Fig. 2b, c and Extended Data Fig. 2e). In five example tomograms, we found no difference between the density of ribosomes inside (cargo) and outside (cytosolic) of autophagic structures (Extended Data Fig. 2f). Next, we compared the nearest neighbour distances (NNDs) of cargo and cytosolic ribosomes in 77 tomograms. Also in this case, neither mean nor median NNDs were significantly different for cargo and cytosolic ribosomes (Fig. 2d and Extended Data Fig. 2g). This suggests that under starvation conditions, autophagosomes mostly engulf cytosol non-specifically and can form without any cargo guiding the membrane.

**Fig. 2:**
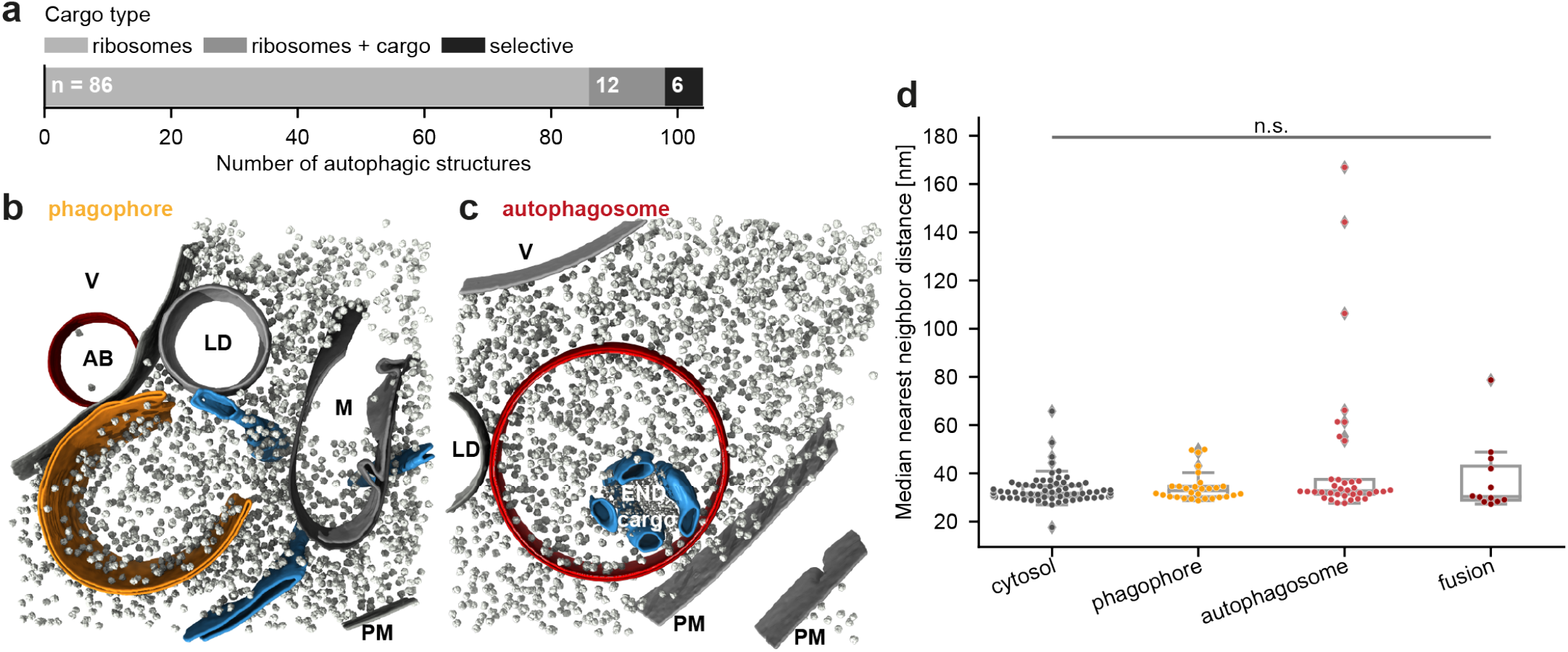
Cargo templating is not essential for autophagosome formation under bulk autophagy conditions. **a**, Numbers of captured autophagic structures containing only ribosomes, ribosomes and selective cargo or exclusively selective cargo. **b**, 3D rendering of an open phagophore (orange) in its native environment, surrounded by ribosomes (grey) and organelles (ER (blue), AB = autophagic body, LD = lipid droplet, M = mitochondrion, PM = plasma membrane). **c**, Segmentation of a closed autophagosome (red) containing ribosomes and the amorphous Ede1-dependant endocytic protein deposit (END) cargo, surrounded by ER (blue). LD = lipid droplet, PM = plasma membrane. **d**, Comparison of ribosome nearest neighbor distances in different compartments. Each dot represents the median distance measured for one tomogram. Differences between cytosol and autophagic ribosome distances were analyzed with the Wilcoxon signed-rank test, treating values from compartments in the same tomogram as paired (n=77 tomograms).

### Phagophores show distinct contact sites with other organelles

Phagophore contacts with other organelles like the ER or the vacuole are known to be crucial for autophagosome biogenesis^5^. To systematically map the subcellular environment of autophagy, we measured the frequency and distance of organelles observed near autophagosomes and phagophores in the tomograms. Particularly the vacuole, ER, nuclear membrane, vesicles and lipid droplets (LDs) were frequently found close to autophagic structures (often < 100 nm, Fig. 3a, b). To confirm these findings, we analysed the colocalization of Atg8-positive structures with other organelles by fluorescence microscopy (Fig. 3c). In agreement with the EM analysis, Atg8 puncta were frequently found at the vacuole (63% of Atg8 puncta) and the ER (61%), specifically at ER exit sites^29^ (ERES, 47%). Unlike mitochondria (<10%), Atg8 puncta still colocalized with LDs in 30% of the cells (Fig. 3c).

**Fig. 3:**
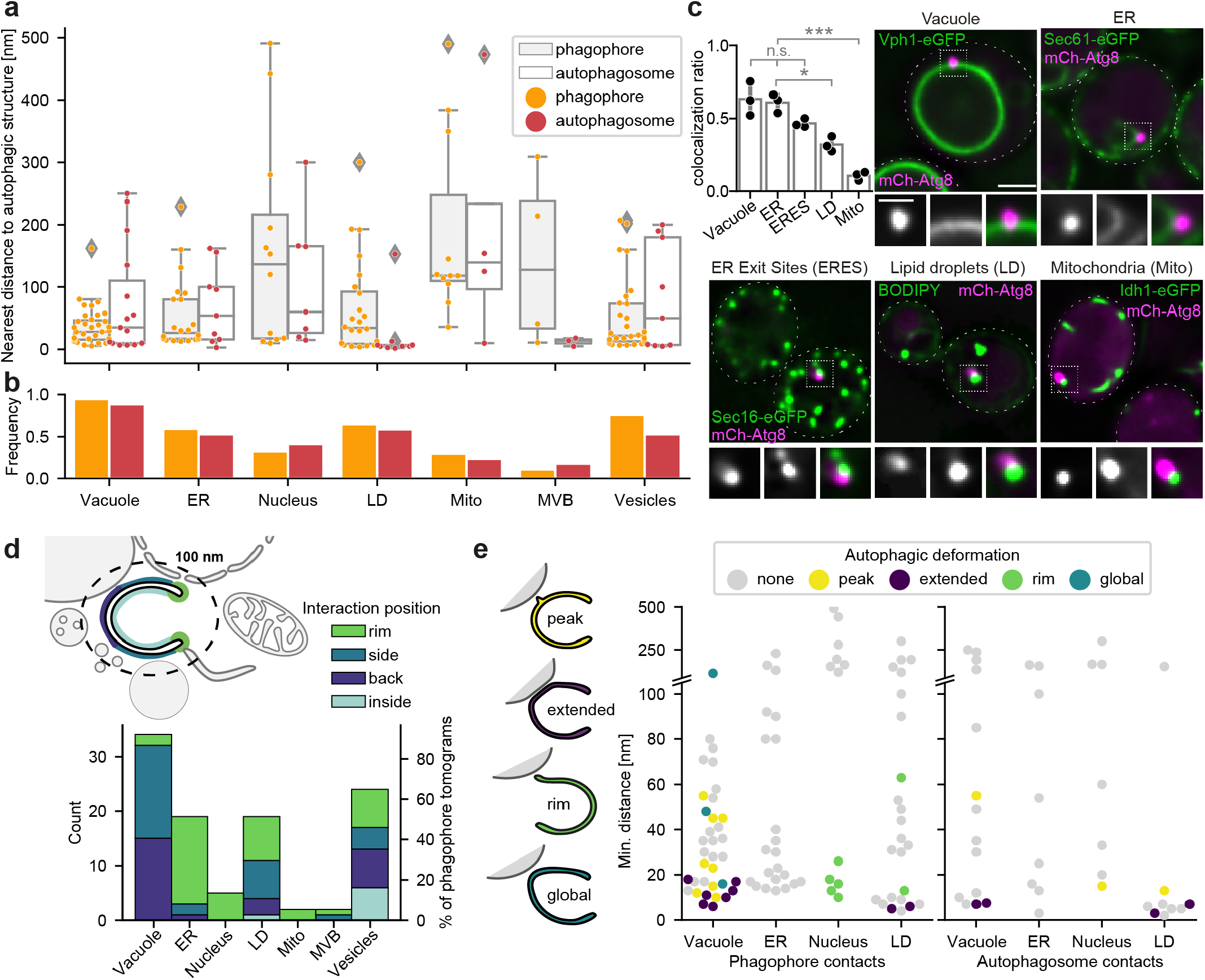
Autophagic structures interact with other organelles. **a**, Nearest distance of different organelles to phagophores (orange) or autophagosomes (red) measured in the tomograms. **b**, Frequency at which different organelles are observed in tomograms of phagophores (n=35) or autophagosomes (n=17). **c**, Quantification and examples of mCherry-Atg8 colocalization with different organelles. For each organelle, the colocalization ratio with Atg8 was measured in three replicates with >100 cells each. **d**, Preferred interaction of organelles with different parts of the phagophore. Contact positions (rim, side, back, inside) are quantified for organelles within 100 nm of the phagophore. **e**, Deformations of autophagic structures at contact sites with other organelles, plotted against the minimum distance for phagophores (left panel) and autophagosomes (right panel). Left, schematic depictions of deformation categories; peaks (yellow), extended contacts (purple), rim deformations (green) and global deformations (petrol).

To distinguish functional contact sites from random ones, we identified two ultrastructural features: First, open phagophores are distinctly polar, with the highly curved rim connecting the inner to the outer membrane. We reasoned that frequent contacts to a distinct part of the phagophore would be a strong indication of a functionally relevant interaction. Accordingly, we assigned each contact with a minimum distance of 100 nm or less to one of the categories “rim”, “inside”, “back” and “side”, based on the area where the closest interaction was observed (Fig. 3d and Extended Data Fig. 3a). Second, in the absence of external forces, a phagophore is expected to adopt a cup-shape form, with the circular rim region as the only high-curvature area^14^. Membrane deformations at contact sites could therefore indicate a specific interaction as they imply additional forces. For quantification, we sorted them into four categories: (1) high-curvature peaks of the autophagic membrane towards another organelle, (2) extended contacts over a large area, (3) extensions of the phagophore rim towards the other organelle (Extended Data Fig. 3b), and (4) global deformations of the whole structure towards the contact (Fig. 3e). Applying these features to the cryo-ET data, four organelles stand out: the vacuole, lipid droplets, the nuclear membrane and the ER.

### The back or side of growing phagophores is anchored to the vacuole

Even though the phagophore assembly site has long been known to localize to the vacuole^7,30^, the tomograms reveal previously unreported aspects of this interaction (Fig. 3, 4a-b). First, open phagophores almost never interact with the vacuole through the rim (Fig. 3d). This leads to a more frequent orientation of the phagophore opening parallel to or away from the vacuole (Extended Data Fig. 3c). Second, half of the phagophores within 100 nm of the vacuole exhibit deformations at the contact sites, with most structures either forming a peak (n = 8) or following the vacuole over an extended area (n = 7) (Fig. 3e, Fig. 4a, b and Extended Data Fig. 3d, e). In contrast, closed autophagosomes show only two extended contacts and one peak in a total of 14 tomograms (Fig. 3e, only wild-type strains). This indicates that these distortions are a characteristic feature of growing phagophores anchored to the vacuole which is largely absent in mature autophagosomes.

**Fig. 4:**
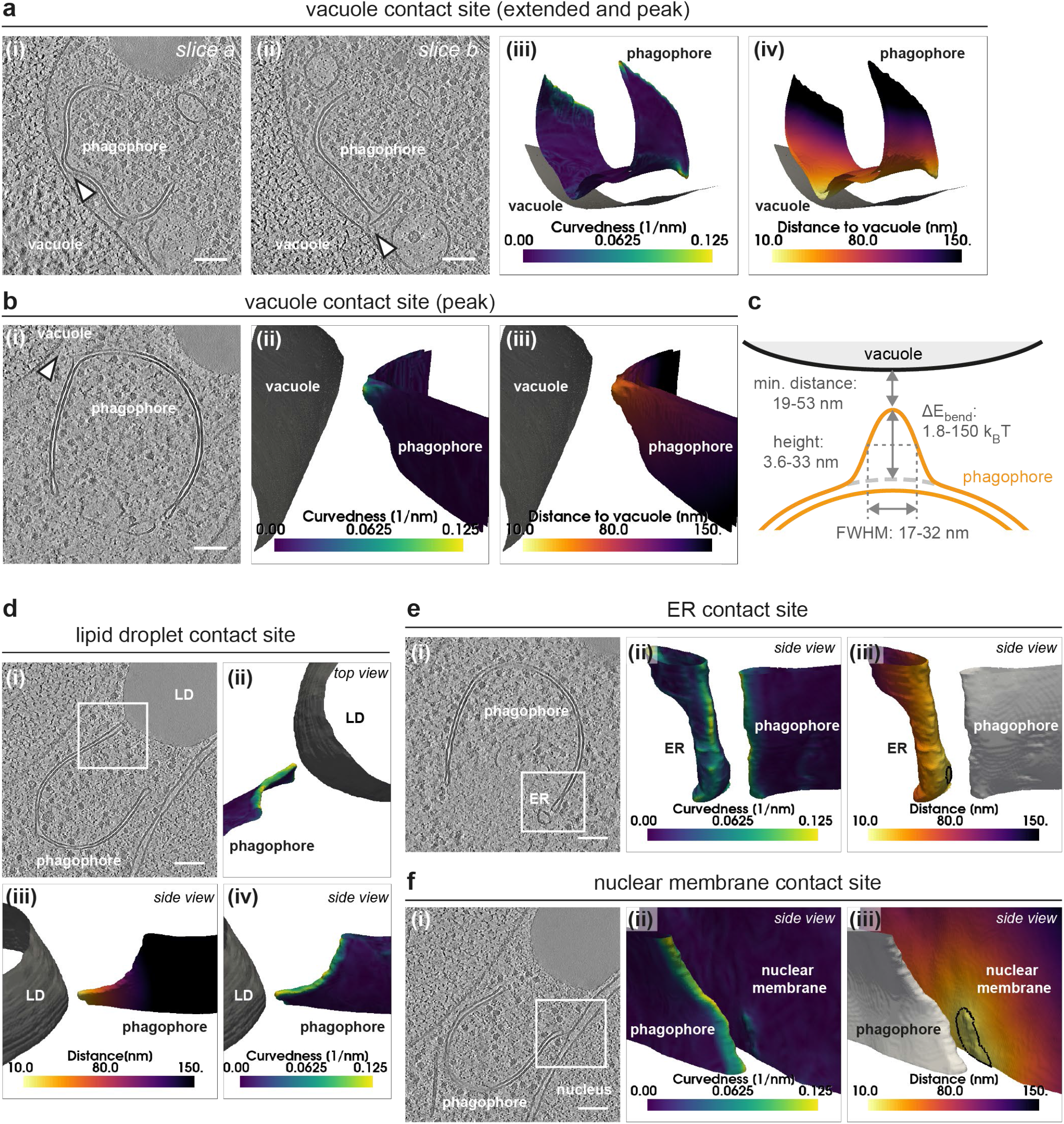
Phagophores engage in specific contacts with the vacuole, lipid droplets, the ER, and nuclear membrane. **a**, Phagophore with a peak and an extended vacuole contact site. (i-ii) Tomogram slices with arrowheads indicating the extended (i) and peak (ii) contact site. Scale bar 100 nm. (iii-iv) Membrane segmentations colored by local curvedness (iii) and distance to the vacuole (iv). **b**, Phagophore forming a peak towards the vacuole. Tomogram slice (i) and segmentations colored by local curvedness (ii) and distance to the vacuole (iii). **c**, Shape parameters (full range) of phagophore peaks towards the vacuole measured for seven peaks. **d**, A rare example of a rim deformation towards a lipid droplet. Tomogram slice (i), top and side views (ii-iv). **e-f**, Phagophore contacts with the ER (**e**) and nuclear membrane (**f**). Tomogram slices (i) and membrane segmentations colored by local curvedness (ii) and distance to the phagophore (iii). The black line on the segmentations marks the membrane areas close enough for Atg2 to bridge the distance.

A detailed analysis of the peak-shaped phagophore-vacuole contact sites reveals that they are highly heterogeneous, with peak heights ranging from 3.6-33 nm, peak widths from 16-32 nm (FWHM), and the minimum distance to the vacuole from 19-53 nm (Fig. 4a-c). The average Pearson’s correlation coefficient between peak elevation and vacuole distance is −0.75, indicating that the phagophore membranes indeed extend towards the vacuole (Extended Data Table 1 and Extended Data Fig. 3f). Extended phagophore-vacuole contacts are equally heterogeneous. Their minimum distances range from 4-20 nm and contact areas from 100-400 nm^2^ (Extended Data Fig. 4g). Thus, apart from the preference for the phagophore side and back, there is no clear consensus structure of the contacts with the vacuole. The difference in phagophore-vacuole spacing also argues against a rigid spacer that would keep the membranes at a fixed distance. However, the local high-curvature areas and deformations in the open phagophores imply that they are physically tethered to the vacuole, withstanding forces strong enough to cause such drastic membrane deformations.

### Lipid droplets associate with autophagic structures and deform phagophores

Lipid droplets are necessary for autophagy in yeast, as their absence inhibits the formation of autophagic structures^31^. LDs have been proposed to act as additional source of lipids for phagophore growth^32^ and as regulators of autophagy by contributing to ER homeostasis as well as maintaining the phospholipid composition^33^. Still, contact sites between LDs and autophagic structures remain largely unexplored^32^. In the tomograms, lipid droplets are found sometimes inside (Extended Data Fig. 2a, b) and often next to both phagophores and autophagosomes (Fig. 3a, 4d) but do not have a preferred phagophore interaction region (Fig. 3d). However, membrane deformations at contacts are observed in two cases in which the phagophore rim clearly extends towards a lipid droplet (Fig. 3e, 4d). While the phagophore-LD distance is rather large in the first case (60 nm, Fig. 3e), the rim gets very close to the lipid droplet in the second (12 nm, Fig. 3e, 4d), thereby suggesting a rare but functional contact.

### The phagophore rim is tethered to the ER and the nuclear membrane

Fluorescence microscopy studies have shown a frequent colocalization of PAS and ER, but also with the nuclear membrane (NM)^29^ (Fig. 3b). This is consistent with the cryo-ET data where tubular ER is observed within 100 nm of the phagophore in more than 50% of the cases (Fig. 3d). Nuclear membrane contacts are rarer, but both NM and ER contacts with the phagophore show a strong preference for the rim (Fig. 3d).

While no strong deformations of the phagophore are observed close to the ER (Fig. 3e, 4e), in all five cases in which the nuclear membrane is within 100 nm of the phagophore, the contact happens through a deformation at the rim towards the nucleus (Fig. 3e, 4f and Extended Data Fig. 4a). Interestingly, phagophore-nucleus contacts cannot only deform the phagophore, but also the nuclear membrane (Fig. 4f), suggesting a strong physical connection between the two organelles. The absence of obvious membrane distortions at contact sites with tubular ER may be explained by its higher motility and lack of physical constraints.

The frequent observation of ER-rim contacts is consistent with their predicted role as lipid transfer sites for phagophore expansion^34^. Recent studies provide strong evidence that lipids are shuttled from the ER to the phagophore through the Atg2/Atg18 complex^12^, known to localize to the phagophore rim^3,29^. Based on the structure of its human homolog^35^, Atg2 could span roughly 20 nm^36^ and phagophores are observed within that distance of the ER or NM in 15 tomograms. In ten of these sites that were analysed in more detail, the ER contact area ranges from 320 to 14500 nm^2^, with a median of around 1600 nm^2^ (Extended Data Table 2). The ER shows a local increase in curvature with decreasing distance to the phagophore in 4/10 tomograms (Extended Data Fig. 4b). In some cases, the phagophore rim and ER are clearly connected by densities with lengths of 17 ± 3 nm (n = 11, Extended Data Fig. 4c), which are however too heterogeneous and rare to be analysed by subtomogram averaging. Still, we note that at all sites with such densities, the local curvature of the phagophore membrane is higher than the local ER curvature (Extended Data Fig. 4d), which might have implications for lipid transfer.

In summary, phagophore-ER contacts almost exclusively happen through the rim and connect the phagophore to the ER or nuclear membrane. The observed membrane deformations suggest a physical connection between the organelles. Based on the short distance and several clearly visible connecting densities, these contacts could function as Atg2-mediated lipid transfer sites.

### Unique structural features of autophagic membranes

To gain detailed insights into the membrane transformations in autophagy, we first identified suitable parameters and developed methods to characterize autophagic structures with minimal manual intervention. Accordingly, membranes were segmented automatically to ensure objectivity of the results^37^. The overall dimensions of the autophagic structures were then estimated from the ~150 nm thick lamella slices by fitting ellipsoids to the inner membranes, and a sphericity index^38^ was calculated for each of the structures (Fig. 5a, b). As estimated from the volumes of the best-fitting ellipsoids (Fig. 5a), phagophores and autophagosomes are overall similar in size. However, while closed autophagosomes are almost perfectly spherical, phagophores show significantly lower sphericity indices (Fig. 5b and Extended Data Fig. 5a, b). This is consistent with the reported elongation of growing phagophores in mouse embryonic fibroblasts^14,39^.

**Fig. 5:**
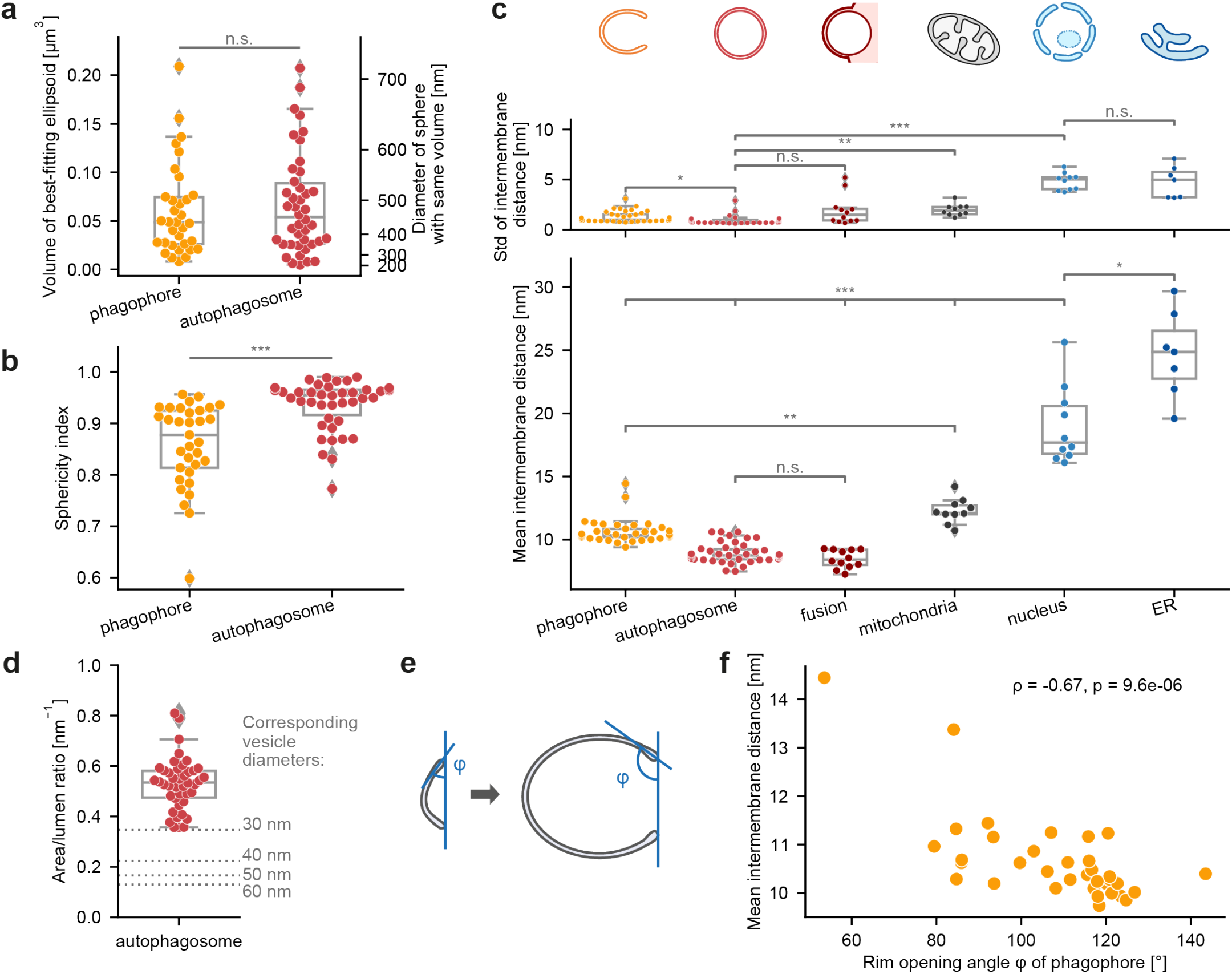
Unique structural features of autophagic membranes. **a,** Overall size of phagophores and autophagosomes estimated by the volumes of the best-fitting ellipsoids. The right axis indicates diameters of spheres with the same volume. **b**, Sphericity index of phagophores and autophagosomes calculated from the best-fitting ellipsoids as 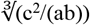 with the ellipsoid axes a>b>c. **c,** Intermembrane distance standard deviations (upper panel) and mean (lower panel) of various double-membrane organelles in the tomograms. Distances were calculated membrane middle to membrane middle; each point represents one structure. **d**, Membrane area to intermembrane lumen ratio of closed autophagosomes. Gray dotted lines show the area/volume ratio of single membrane vesicles with indicated diameters. **e**, Scheme showing the rim opening angle φ for two phagophores in different stages of growth. **f**, Intermembrane distance plotted against φ. The mean intermembrane distance correlates with the rim opening angle with a Spearman’s correlation coefficient of −0.67, p = 9.6·10^-6^. Statistical analysis: a & b: Mann-Whitney-U test; c: Kruskal-Wallis H-test and pairwise Games-Howell post-hoc test. Autophagosomes n=42, phagophores n=37, fusion n=12, mitochondria n=10, nucleus n=10, ER n=7. ***p<0.001, **p<0.01, *p<0.05, n.s.: p≥0.05.

A crucial parameter that determines the shape of autophagic structures is the distance between the inner and outer membrane. To quantify this intermembrane spacing, we developed a robust minimum distance algorithm (Supplementary Note 1), which can handle holes and overhangs in segmentations and measures the distance from thousands of points. Notably, the average intermembrane distance of autophagic structures is significantly smaller than in mitochondria, the nuclear membrane, and ER sheets (Fig. 5c). The observed 9-11 nm spacing (measured from middle of one phospholipid bilayer to middle of the other bilayer) is also smaller than previously reported for autophagosomes (20-50 nm in general, < 30 nm in yeast)^40^. Both autophagosomes and their fusion intermediates display similar values (8.9 ± 0.79 nm and 8.5 ± 0.72 nm, mean ± SD, n = 42 and n = 12, respectively) which is strikingly homogenous across the whole membrane (Fig. 5c). This makes the intermembrane distance of autophagic structures a unique feature to distinguish autophagosomes from other structures in the cell.

Because autophagosomes are close to perfect double-membrane spheres, their full structures can be extrapolated based on tomogram data. This allows us to estimate for each observed autophagosome its total membrane area to intermembrane lumen ratio, yielding an average area/lumen ratio of 0.53 ± 0.10 nm^-1^ for closed autophagosomes (Fig. 5d). Of the two major processes thought to sustain phagophore growth, (1) vesicle fusion^41^ and (2) direct lipid transfer^42^, only the first adds volume to the intermembrane lumen. The reported size range of vesicles contributing to phagophore growth^9,4^ is 30-60 nm for Atg9 and > 60 nm for COPII^43^, corresponding to a membrane area/volume ratio of 0.35-0.13 nm^-1^ (grey dotted lines, Fig. 5d). To define the *in situ* vesicle landscape, we measured the average diameter of vesicles observed within 100 nm of phagophores. On average those vesicles have a diameter of 40 nm (Extended Data Fig 5d, e), which fits the expected range for Atg9 vesicles. By comparing the area/volume ratios of vesicles and autophagosomes (Fig 5d), it is clear that if the intermembrane lumen of autophagosomes is built from vesicles alone, they do not contribute enough membrane to build the whole autophagosome, arguing for lipid transfer from the ER as a major membrane source during phagophore expansion.

Assuming that the intermembrane lumen of autophagosomes does not change by other means, our data suggest that between 60 - 80% of the membrane area is derived from lipid transfer or synthesis (Extended Data Fig. 5f).

Interestingly, compared to closed autophagosomes, open phagophores show a significantly higher mean intermembrane spacing (10.6 ± 0.93 nm, n = 37, Mann-Whitney-U test p = 2.2·10^-11^). This suggests that counter to previous assumptions^14,44^, the intermembrane distance of phagophores is not constant, but rather decreases during expansion. Testing this hypothesis required a method for sorting the phagophores by degree of maturation. Having evaluated different parameters (Supplementary Note 2), we found the rim opening angle φ, calculated as the mean angle between a plane through the rim opening and tangent planes to the inner phagophore membrane at the rim, to be the most indicative (Fig. 5e). Throughout phagophore growth, φ should increase from around 0° in the initial membrane disk to 180° just before closure into a double-membrane sphere. As hypothesized, the mean intermembrane distance of the captured phagophores decreases significantly with φ (Fig. 5f, Extended Data Fig. 5g-i). Taken together, the analysis yields conclusive evidence that the intermembrane distance decreases during autophagosome formation.

### The phagophore rim shape transforms during phagophore growth

A striking feature of phagophores over autophagosomes is the highly curved rim at the opening of the cup-shaped structure. Notably, upon inspection of the tomograms the rims of many phagophores appeared dilated (Fig. 6a). This is in contrast to the half toroid shape assumed in the literature^14,44,45^, suggesting a direct impact on the rim curvature and bending energy.To further investigate this phenomenon, we produced refined segmentations of 26 well-resolved phagophore rims and used custom scripts to detect their tips (Extended Data Fig. 6a-d). The distances between the inner and outer membrane were measured orthogonally to the rim direction and mapped against their distance from the phagophore tip (Extended Data Fig. 6d). Plotting for each rim the mean intermembrane spacing against the distance from the tip (Fig. 6b) suggests that all analysed rims show swelling. To determine if indeed each rim is dilated, we next checked for the presence of maximum and minimum peaks in the intermembrane distance along each segmented rim, as well as their height and distance from the tip (Extended Data Fig. 6e, Extended Data Table 3). The peak analysis confirms that all rims show a clear intermembrane distance maximum when moving from the tip towards the back, and maxima are present along the complete rim segment in most examples (Extended Data Fig. 6e). In contrast, minimum peaks, i.e. a constriction of the membranes after the swelling, are observed less consistently and therefore not analysed in more detail. The position of the dilation maximum differs substantially between rims (17 ± 7 nm from the tip) (Fig. 6b, c) and within the individual structures (median standard deviation 2.6 nm). The intermembrane distances are less variable, with a mean maximum distance of 14.7 ± 1.8 nm and an average 10.9 ± 0.52 nm “base” distance for the back part of the rims excluding the dilated region. By dividing the respective maximum and base distances, a “dilation factor” is obtained for each rim, with a mean value of 1.35 ± 0.15 (±SD) (Fig. 6c).

**Fig. 6:**
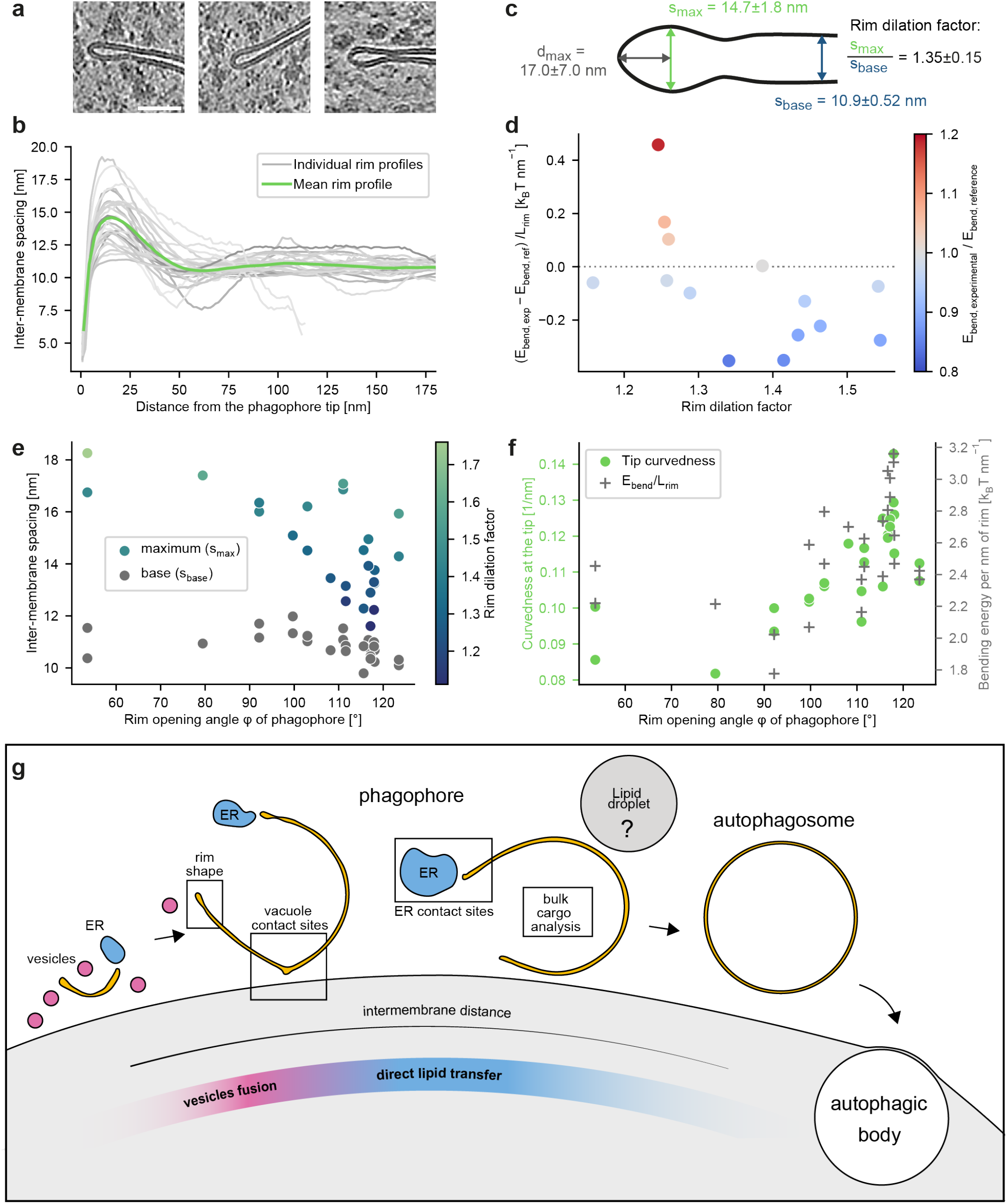
Characterization of the phagophore rim and model of autophagosome biogenesis. **a**, Example tomogram snapshots of phagophore rims, scale bar: 50 nm. **b**, Rim shape parameters including intermembrane spacing in the back (s_back_) and at the swelling maximum (s_max_), and the distance of the maximum to the tip (d_max_). Mean and standard deviation calculated from 26 rims. **c**, Mean (green) and individual rim profiles (gray) for all analyzed rims (n=26), plotted as intermembrane distance vs distance from the tip. **d**, Effect of rim swelling on the bending energy of experimental (E_bend, exp_) vs. hypothetical non-swollen reference rims (E_bend, ref_) (n=14). The experimental is smaller than the reference bending energy for most cases with a dilation factor (s_max_/s_back_) of 1.3 or higher. **e**, Rim intermembrane spacing (maximum and back) vs. rim opening angle φ. Spearman’s rank correlation: ρ = −0.64, p = 4.1·10^-4^ (max. spacing), ρ = −0.63, p = 5.4·10^-4^ (back spacing), ρ = −0.44, p = 0.025 (dilation factor) (n=26). **f**, Membrane curvedness within 1.408 nm (1 binned pixel) of the tip (green, left y axis) and bending energy per nm of rim (grey, right y axis) plotted against φ. Spearman’s rank correlation: ρ = 0.77, p = 4.1·10^-6^ (curvedness), ρ = 0.53, p = 0.0057 (bending energy) (n=26). **g**, Summary of ultrastructural features of autophagy and autophagosome biogenesis model: vesicles (magenta) are major contributors during early biogenesis, while direct lipid transfer from the ER (blue) is the main source for membrane growth in later stages. This results in a decrease of the intermembrane spacing and increase of rim curvature, favoring rim constriction towards the closed autophagosome.

We speculated that the observed rim swelling might reduce the local mean curvature and thus the bending energy compared to a non-dilated structure. By widening, the rim locally approaches the zero-energy catenoid shape with principal curvatures of equal magnitude but opposite sign. To test this hypothesis, we constructed artificial, non-dilated versions of the analysed rims, keeping the same membrane area, overall shape and base intermembrane distance (Extended Data Fig. 6f). Figure 5d shows the difference between the respective Helfrich bending energies^46^ for the experimental and reference rims, normalized by the length of the rim segments and plotted against the dilation factor. While no clear trend is observed at dilation factors below 1.3, all experimental rims with a dilation factor > 1.3 show a lower (n = 7 rims) or equal (n = 1 rim) bending energy compared to their non-dilated counterparts. This suggests that strong swelling indeed decreases the bending energy at the rim, which helps stabilizing the open phagophore state.

How does the shape of the rim evolve with phagophore growth? The maximum intermembrane spacing decreases strongly with the rim opening angle φ (Fig. 6e and Extended Data Fig. 6g). As a result, both the curvature at the rim tip and the bending energy per nm of rim increase during phagophore growth (Fig. 6f, Extended Data Fig. 6h, Extended Data Table 4). Interestingly, these dynamic changes appear to have two independent and additive causes: First, a decrease in rim dilation (Fig. 6e) consistent with approaching locally a catenoid shape upon tightening of the neck, which decreases the energetic cost of a high first principal curvature. Second, however, the base distance also decreases with phagophore growth (Fig. 6e), which is most likely not a consequence but rather a driver of rim constriction. In this model, the decrease of phagophore membrane spacing (1) increases the first principal curvature at the rim and therefore (2) favours rim constriction to reduce the rim length and minimize the overall bending energy, thus promoting phagophore closure.

## Discussion

Our structural analysis of autophagy *in situ* shows that phagophores are unique organelles that engulf mostly bulk cargo under starvation and form distinct contacts to the vacuole, ER and rarely to lipid droplets. Unexpectedly, their already thin intermembrane spacing decreases even more during growth, concomitant with a gradual decrease of rim swelling and increase of the rim curvature (Fig. 6g).

From the structures of closed autophagosomes, we estimate that only 20-40% of their membrane is contributed by fusion of vesicles. Note that 35-135 vesicles (60-40 nm diameter) would suffice to build the intermembrane lumen of a typical autophagosome and that no processes have been described to date that actively reduce the intermembrane lumen of phagophores. Lipid transfer should then contribute 60-80% of the autophagosome membrane, supported by the frequently observed close contacts of the phagophore rim with the ER and NM. Even if the luminal volume expands slightly to counteract the high rim curvature and tight membrane spacing, this will only decrease the number of needed vesicles and necessitate even more lipid transfer (Extended Data Fig. 5j).

In yeast, Atg9 vesicles contribute mainly to the initial nucleation stage^8^, whereas other membrane sources such as COPII vesicles are thought to contribute to both early and late phagophore growth^11^. The decreasing intermembrane distance from early to late phagophores implies that their area/lumen ratio is smaller initially and increases as they grow. This is in line with a model in which vesicles mainly contribute to the initial stages, whereas lipid transfer becomes the major membrane source later during phagophore growth (Fig. 6g).

How does the proposed shift in membrane sources affect phagophore growth? In the absence of other mechanisms controlling the luminal volume, the relative rates of vesicle fusion and lipid transfer determine the rate of phagophore thinning. Phagophore thinning increases in turn the curvature at the rim, which should accelerate constriction towards an almost closed phagophore. Thus, we speculate that the size of the final autophagosome might be limited by the fusion rate and total number of contributing vesicles. How the recruitment of the ESCRT machinery for phagophore closure^47^ relates to the maturation of the rim and to the final autophagosome size remains to be determined.

Our analysis of the organelle interactome of autophagic structures shows that phagophores form very polarized contact sites. The most prominent is the ER-rim contact site, which likely functions in Atg2-mediated lipid transfer^12^. Based on fluorescence microscopy experiments, Atg2 localizes to the phagophore rim^3,29^. In agreement with this, we identified electron dense structures spanning across the ER-rim contacts with lengths in the expected range for Atg2^36^ (Extended Data Fig. 4c). Calculations based on our analysis suggests that less than 100 copies of Atg2 per phagophore would suffice to build an average-sized autophagosome (Supplementary Note 3). This could explain why so few connecting densities are found at the contact sites.

Even more frequently than phagophore-ER contacts we observe interactions with the vacuole, preferentially at the back or side of the phagophore. The heterogeneity of these contacts is consistent with a recent study showing that the anchoring of the PAS machinery to the vacuole is an avidity-driven process mediated by locally clustered Vac8^48^. If the tethering of the phagophore membrane is also mediated by Vac8, variable cluster sizes could explain the variety of contact shapes observed. As to why phagophores are tethered to the vacuole, we suggest that this arrangement allows the starving cell to produce many autophagosomes one after the other while keeping the autophagy machinery at a defined location for growth, maturation and fusion. In fact, we often observe partially enveloped autophagic bodies in the vacuole directly next to open phagophores (Fig. 1d and Extended Data Fig. 2d), suggesting that the same site was used at least twice in quick succession.

Finally, the present study can serve as blueprint for studying organelle biogenesis processes. Specifically, induction of organelle formation, fluorescent tagging of the growing structure, and genetic or pharmacological manipulation to accumulate intermediates are essential steps to capture the intermediates by correlative cryo-ET. Although it is static by nature, *in situ* tomography is not limited to resolving protein structures, but can also provide information on the dynamic morphology of membranes and direct measurements for the biophysical characterization and modelling of transient cellular processes.

## Methods

### Yeast strains

A list of budding yeast (*S. cerevisiae)* strains used in this study is provided in Supplementary Table 1. All of the yeast strains were based on the DF5 background. Standard protocols for transformation, mating, sporulation and tetrad dissection were used for yeast manipulations^49^. Chromosomally tagged strains and knockout strains were constructed using a PCR-based integration strategy^50^. Standard cloning techniques were used.

### Live-cell fluorescence microscopy

For fluorescence microscopy, yeast cells were grown in synthetic growth medium supplemented with all essential amino acids and 2% glucose. The next day, cells were diluted to OD 0.1 and grown until mid-log phase (0.5–0.8 OD) before imaging. Microscopy slides were pretreated with 1 mg ml^-1^ concanavalin A solution. Widefield imaging was performed at the Imaging Facility of the Max Planck Institute of Biophysics using a Nikon Ti2 Eclipse microscope comprising, an Olympus Apo TIRF 100x 1.49 oil objective and a Hamamatsu ORCA-Flash 4.0 LT+ Digital CMOS camera. The images were deconvolved using the Nikon NIS Elements Batch Deconvolution Tool (automatic function). Image analysis was performed using the CellCounter plugin in ImageJ (https://imagej.nih.gov/ij/).

### Cryo-ET sample preparation

A detailed protocol of the correlative cryo-ET is available under dx.doi.org/10.17504/protocols.io.e6nvwkz4wvmk/v1.

#### Starvation & plunge freezing

Yeast cultures were inoculated from overnight cultures in YPD medium (1% yeast extract, 2% peptone and 2% glucose) to an OD600 of 0.15 and grown at 30°C to an OD600 of 0.8. At this point, medium was switched to SD-N (synthetic minimal medium lacking nitrogen; 0.17% YNB without amino acids and ammonium sulfate, supplemented with 2% glucose) and cells were incubated for a time span of 0.5-3 hours at 30°C. For 3D-correlation on the grid, 1 μm Dynabeads (Dynabeads MyOne Carboxylic acid #65011, Thermo Fisher Scientific) were added to the cells at a dilution of 1:20. Grids (200 Mesh Cu SiO2 R1/4, Quantifoil) were plasma cleaned for 30 s before plunging. 4 μl of starved cell solution with beads was applied on the grid, blotted and plunged in ethane-propane with a Vitrobot Mark IV (Settings: Blotforce = 8, Blottime = 10s, room temperature).

#### Cryo-CLEM and FIB milling

Grids were mounted on modified autogrids with cut-out for FIB-milling and fluorescence image stacks were acquired on a cryo-confocal microscope (Leica SP8 with Cryo-Stage) equipped with a 50X/0.9 NA objective (Leica Objective #506520) and two HyD detectors. Stacks (Step size 300 nm, x-y pixel size 85 nm) were acquired using 488 nm and 552 nm laser excitation for eGFP- and mCherry-labelled proteins, respectively. In the case of eGFP only strains (eGFP-Atg8 and eGFP-Ede1/Δypt7), signal from autofluorescent Dynabeads was acquired as second channel corresponding to red emission wavelengths to easily distinguish fiducial beads from cellular signal. Stacks were deconvolved using Huygens (Scientific Volume Imaging, http://svi.nl). Target sites corresponding to Atg8 puncta or Ede1 END cargo were 3D-correlated to SEM/IB images in the FIB/SEM microscope (FIB Scios and Aquilos, Thermo Fisher Scientific) using the 3D-Correlation Toolbox (3DCT)^20^. Lamellae were milled in correlated sites as described in a previously published protocol^21^. In a few cases (e.g Extended Data Fig. 1a), a widefield microscope integrated in the FIB/SEM chamber (METEOR, delmic) was used to confirm the presence of fluorescence signal in the lamella, as previously published^51^.

#### Cryo-EM data Acquisition

Tomograms were acquired on a Transmission Electron Microscope (Titan Krios, FEG 300kV, Thermo Fisher Scientific) equipped with an energy filter (Quantum K2, Gatan) and a direct detection camera (K2 Summit, Gatan) at a magnification of 42000x (pixel size 3.52 Å) and defocus ranging from −5 to −3.5 μm. Positions for tomogram acquisition were determined by correlation of fluorescence data to TEM images of the grid squares containing lamellas (3DCT), followed by inspection of low-magnification lamella images. Frames were recorded in dose-fractionation mode, with a total dose of 120 e^−^/A^2^ per tilt series using SerialEM^52^. A dose-symmetric tilt scheme was used with an increment of 2° in a total range of ±60° from a starting angle of 10° (+ or −) to compensate for lamella pre-tilt (mostly around 11°). Frames were aligned using MotionCorr2^53^ and reconstruction was performed in IMOD by using the TomoMAN wrapper scripts^54^.

#### Detection & averaging of ribosomes

Ribosome positions were determined by template matching on 2x binned tomograms (IMOD bin 4, 14.08 Å pixel size) using the StopGAP^55^ software package. In brief, a reference was constructed from ~300 manually picked ribosomes, which were aligned in StopGAP and used for template matching. For each tomogram, the positions of the 1500 highest scoring peaks were extracted and saved. Tomograms were exported from TomoMAN to Warp/M^56^ for CTF estimation and locally reconstructing particles. Classification and refinement of 1x binned subtomograms (pixel size 7.04 Å) was performed in Relion 3.1.2^57^, which yielded the final list of particles and a ribosome structure at 15.1 Å resolution (0.143 FSC criterion, Extended Data Fig. 2e).

#### Segmentation and visualization

Tomograms at 2x binning with a nominal pixel size of 1.408 nm were denoised using cryo-CARE on tomograms reconstructed from odd/even frames^58^. Membrane middles (middle of phospholipid bilayer) were detected automatically using TomoSegMemTV^37^ and selected in Amira (Thermo Fisher Scientific). Segmentations for display purposes (Fig. 1i-l) were manually refined in Amira, gaussian filtered and displayed in ChimeraX^59^. For analyses of membrane curvature (phagophore rims and contact sites), the automatic segmentations were refined manually in Amira. Mesh generation from the filled segmentation and curvature determination was done using PyCurv^60^ using a radius hit of 8 nm. Visualizations of different parameters on segmented membranes (Fig. 4 and Extended Data Fig. 3d-e, 4a) were produced with Pyvista^61^.

### Data analysis and statistics

All analyses of membrane structures and particle distributions were performed with custom written scripts in Python 3. Major python packages used in this work include numpy^62^, scipy^63^, pandas^64^, Pyvista^61^, scikit-learn^65^ (data analysis), matplotlib^66^, seaborn^67^, Pyvista (visualization), tifffile^68^, mrcfile^69^, starfile^70^ (data I/O). Statistical analyses were performed with the statistical analysis package in scipy (scipy.stats) and the pingouin package^71^, using the tests indicated in each respective analysis. In general, statistical analysis of differences between two groups was performed using the Mann-Whitney *U* test for independent and the Wilcoxon signed-rank test for dependent samples. Comparison of more groups was performed with the Kruskal-Wallis H test and pairwise Games-Howell post-hoc test. Correlation between variables was assessed with Spearman’s rank correlation coefficient.

### Ribosome density and nearest neighbour analysis

For ribosome density estimations, we produced complete filled segmentations of all organelles, large structures like glycogen granules and the volume outside the lamella in 5 example tomograms at 2x binning using Amira. For tomograms of autophagosomes and autophagic bodies, the autophagic content was segmented directly and the initial cytosol voxels were defined as all unlabelled voxels. For phagophores, a convex hull around the phagophore membrane points was calculated and all unlabelled voxels in the hull were subtracted from the cytosol and labelled as initial phagophore points. Next, the initial cytosol and phagophore volumes were cleaned with binary opening in scipy using a 3×3×3 voxel cube as structuring element and 2-3 iterations. The resulting cytosol and autophagic content segmentations were then used to assign ribosomes and calculate ribosome densities in- and outside the autophagic structures. For nearest neighbour analyses, we circumvented the time-consuming full segmentations by directly using TomoSegMemTV-generated segmentations of the autophagic membranes, only deleting parts of the automatic segmentation manually that deviated significantly from the actual membrane position. Convex hulls around the roughly segmented membranes were used to assign ribosomes to autophagic content or cytosol, and nearest neighbour distances within each area were calculated with KDTrees using scipy. Both (1) an unpaired Kruskal-Wallis test of mean and median nearest neighbour distances in cytosol, phagophores, autophagosomes and fusion structures, and (2) a Wilcoxon test treating measurements from the same tomogram as paired measurements revealed no significant differences between the ribosome nearest neighbour distances inside and outside of autophagic structures.

### Contact site analysis

For an analysis of the contact sites of autophagic structures with other organelles, we excluded all tomograms from the *Δypt7* strain since the overall cellular architecture in this strain was disturbed by accumulation of medium-sized vacuoles^72^ (Extended Data Fig. 1i). The nearest distances between different organelles (Fig. 3a) were measured manually in the tomograms using IMOD (estimated precision ~ 2 nm). For the preferred interaction areas, all contacts at the rim up to the dilation maximum were counted as rim contacts, while the back was defined as the area opposite of the rim. The orientation of phagophores relative to the vacuole was assessed by calculating the angle α between the rim plane normal (pointing outwards) and the vector of the phagophore point closest to the vacuole to its nearest point in the vacuole. For the phagophore deformations at contact sites (Fig. 3e), peak contacts were defined by a high curvature and a local increase in the phagophore intermembrane distance, while extended contacts do not show strong changes in the intermembrane distance, but are usually accompanied by a local flattening of the phagophore membrane. Global deformations are defined as a strong deviation from the usual spheroid-like shape of the phagophore towards the other organelle in the absence of a clear peak or extended contact. Finally, rim deformations are clear deviations of the rim tip out of the rim plane and/or the best-fitting rim circle. In an analysis of the maximum distance of rim points from the best-fitting rim plane, these phagophores stand out with high plane distances despite a phagophore orientation in which the rim is clearly visible, ruling out segmentation inaccuracies as cause of the deviation (Extended Data Fig. 3b).

#### Vacuole contact peak analysis

For a detailed analysis of peaks in the phagophore membrane towards the vacuole, we produced refined segmentations of the phagophore membrane at the relevant sites, determined the curvature using PyCurv^60^ (radius hit 8 nm) and calculated for each phagophore point its nearest-neighbour distance to the segmented points of the vacuole. The resulting curvedness and vacuole distance values were visualized in Fig. 4 using Pyvista. Having tested different potential parameters for an automatic detection of the peaks in the outer phagophore membrane, the best-performing parameter was the product of the local gaussian curvature with the distance to the inner membrane. We calculated this value for all points in the outer phagophore membrane and applied a threshold of mean+3*std to get potential peak points. The resulting points were clustered with DBSCAN (ε = 10 nm) and all point clusters further than 10 nm from the segmentation border were included in the following analysis. For each peak, its full area was extracted by applying a cutoff to the distance to the original cluster points, which was determined as the distance at which the first derivative of the intermembrane distance vs distance to cluster points exceeds an empirically determined value (−0.15). The height *h* of each peak was determined from the maximum intermembrane distance *d_max_* and the intermembrane distance of the whole segmented phagophore piece excluding the peak areas, *d_base_,* as *h* = *d_max_ d_base_*. The width (FWHM) of each peak was estimated as the diameter of a circle fit into the points with *d_max_* – *d_base_* ≈ 0.5h. Finally, the bending energy stored in the peak was estimated as difference in Helfrich bending energies ΔE_bend_ = E_bend,peak_ – E_bend,base_, where E_bend,base_ is the bending energy of the base mesh normalized to the same membrane area as the peak area. To allow a better comparison of the different peaks as well as extended phagophore-vacuole contacts, we produced 2D elevation and vacuole distance maps using in-house scripts which are available upon request (Extended Data Fig. 3f, g).

#### ER contact site analysis

For ten ER-phagophore contact sites, the local ER and phagophore membrane segmentations were refined carefully and the curvature was determined using Pycurv (radius hit 8 nm). Assuming a membrane thickness of around 5 nm^73^ and an Atg2 protein length of 20 nm^12^, we applied a cutoff distance of 25 nm to the ER-phagophore distance measured between membrane middles to analyze the size, local curvature (Extended Data Fig. 4a) and curvature change with interorganelle distance (Extended Data Fig. 4b) of potential contact regions. Potential Atg2 densities were identified visually and their start and end coordinates were marked with IMOD. These coordinates were used to estimate the length of the density and find the closest cells in the phagophore and ER membrane meshes to report the local curvatures (Extended Data Fig. 4d).

#### Analysis of membrane morphology of autophagic structures

All autophagic structures containing at least partially cytosolic cargo (visible ribosomes) were included in the following analysis, including structures observed in the eGFP-Ede1 *Δypt7* strain. We defined as phagophores all cytosolic autophagic structures with a visible opening and rim, the rest as autophagosomes. While some phagophores could have been mislabeled as “closed autophagosomes” if the rim was completely outside the lamella volume, the significant morphological differences between the autophagosomes and phagophores e.g. in sphericity and intermembrane distance, together with the fact that autophagosomes show a similar intermembrane distance as the subsequent fusion structures, clearly argues that most autophagosomes were assigned correctly. We used the automatically generated membrane segmentations for this analysis as described above, only deleting parts of the automatic segmentation manually that deviated significantly from the actual membrane position. Note that since the segmentations mark the middle of the membrane bilayer, all distances and fit results are reported with respect to the membrane middles.

#### Analysis of intermembrane distances

To analyse intermembrane distances, we devised a refined minimum distance algorithm that only uses minimum point distances and is robust against peaks and holes in either of the two surfaces. The algorithm is described in detail in Supplementary Note 1. For the comparison of different double-membrane organelles, we segmented cortical and cytosolic ER sheets as well as mitochondrial membranes without cristae. The phagophore membranes were divided at the rim into inner and outer membrane. For the autophagosome-vacuole fusion structures, only the distances in the wrapped part were analyzed. This was done by automatically detecting the points at the border of the wrapped area, fitting a plane through these points and analysing only the membranes on the side of the plane that faces away from the vacuole (Extended Data Fig. 5c).

#### Size and sphericity measurements

To estimate the size and sphericity of whole autophagosomes and phagophores even though only a section of each structure is present in the tomograms, we fit ellipsoids into the inner and outer membrane segmentation points. We used an iterative ellipsoid algorithm described in and adapted from Kovac et al.^74^ which is more robust than simple least-squares approaches. To give a rough estimate of the overall dimensions of the structures, we report the volumes of the best-fitting ellipsoids for the inner membranes of both phagophores and autophagosomes. Since phagophores are incomplete, this volume is not the engulfed volume, but rather reflects roughly the expected final volume. To estimate how spherical the structures are, the best-fitting ellipsoids to the inner membranes were used to calculate the sphericity index as described in Cruz-Matías et al., as 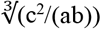 with the ellipsoid axes a>b>c^75^. Additionally, we calculated the “classical” sphericity according to its original definition as ratio between the surface area of a sphere with volume equal to the structure of interest and the structure surface area^76^ (Extended Data Fig. 5b). Finally, we also applied a least-squares algorithm for sphere fitting and used the root mean square error (RMSE) of the sphere fit to the inner membrane as a third parameter for estimating how well phagophores or autophagosomes correspond to a sphere (Extended Data Fig. 5a).

#### Membrane source contribution calculations

The area-to-lumen ratios of autophagosomes were estimated based on the ellipsoid fits to the inner membrane. To ensure that the modelled autophagosome has the correct intermembrane distance, we modelled the outer membrane by adding the mean intermembrane distance determined for the respective autophagosome to all axes of the inner ellipsoid. The surface area of the inner and outer ellipsoid was estimated using the Knud Thomsen approximation^77^ which gives a computationally inexpensive estimation of the ellipsoid area with a maximum error of ±1.061%. Since these ellipsoids correspond to the middles of the inner and outer membrane, we corrected the axis lengths with half the membrane thickness (0.5*5 nm = 2.5 nm, addition to inner and subtraction from outer ellipsoid axes lengths) to calculate the intermembrane lumen. The same correction was performed for calculating the area/volume ratio of vesicles. The diameter of vesicles within a radius of distance of 100 nm from phagophores in the tomograms were measured manually in IMOD.

To calculate the contribution of lipid transfer vs vesicle fusion to the final autophagosome membrane, we assumed that the intermembrane lumen of autophagosomes V_AP_ corresponds to the combined lumen of all fused vesicles V_ves_, and that the membrane area A_AP_ is the sum of the membrane areas of the vesicles A_ves_ and the area of lipids transferred e.g. from the ER, A_ER_, thus:

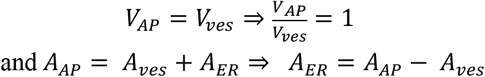

The contribution of lipid transfer to the final membrane, A_ER_/A_AP_, can thus be calculated from the area-to-volume ratios of the vesicles and autophagosome, R_ves_ = A_ves_/V_ves_ and R_AP_ = A_AP_/V_AP_, with:

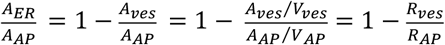

#### Analyzing the completeness of phagophores

One challenge in the analysis of phagophores was to find a parameter to robustly estimate the completeness of each observed structure. A comparison of all considered parameters is given in Supp. Note 2. The most robust parameter that we identified is the “rim opening angle” φ, defined as the angle between a plane through the phagophore rim and tangential planes to the phagophore membrane close to the rim. To measure the angle for each phagophore, a plane was fit through the roughly segmented points at the rim tip to give the rim plane. All inner membrane points within 50 pixels (~70 nm) to the rim plane were used to fit the tangent planes in the following manner: Using a circle fit through the rim points, these inner membrane points were divided into angular batches spanning 10° each. Next, planes were fit to each batch of points and the normals of these tangential planes were used for angle calculation with the rim plane normal. In practice, we noticed that this measurement sometimes leads to high variances of the angle, presumably due to errors in the determination of the rim plane normal or inherent asymmetry of the phagophore. To counteract this, we reasoned that the normals of the rim tangent planes should form a cone whose base plane should ideally be parallel to the rim opening plane. We therefore calculated the cone base planes for all rims in which the available points span more than 90° of the full cone circle, used the normals as corrected rim plane normals and reported the final rim opening angle as the mean of the angles between this vector and all tangential plane normals.

#### Correlation and bootstrap analysis to test for significance

Pair-wise correlation of different parameters was assessed with Spearman’s rank correlation coefficient. However, the input parameters were often mean values of different measurements and the simple correlation analysis did not consider the spread of raw data resulting in these mean values. To take the raw data into account and analyse the confidence of the reported correlation coefficients, we used a bootstrapping approach as described by Curran^78^. In brief, for 10^5^ iterations, we replaced each mean value by a randomly chosen value from its raw distribution and recalculated the Spearman correlation coefficient and p value. The resulting distributions of correlation coefficients and p values are shown in Extended Data Figures 5h & 6g-h.

### Phagophore rim analysis

To analyze the phagophore rim shapes in detail, we produced refined segmentations of 26 rim segments and determined the local membrane curvatures using PyCurv (radius hit 8 nm). We next developed a strategy to extract the tip points and two sides of each rim segment and approximated the general shape of the phagophore at the rim segments by fitting a 3^rd^ order polynomial surface through points lying in the middle between the two sides (see Extended Data Fig. 6a+d). The intermembrane spacing of the rims orthogonally to the overall rim shape was then calculated by ray tracing from one side of the rim to the other using the normals of the closest middle surface points as ray directions. To generate 1D and 2D histograms of distance and curvature values, all rim points were mapped to their nearest middle surface point, and we calculated for each middle surface point (1) its closest geodesic distance along the middle surface to the smoothed tip (“distance from the tip”) and (2) the geodesic distance of its closest tip point to the tip point with the lowest z (“distance along the tip”). Using these values as xy coordinates, we binned the points using 1-pixel bins for the distance from the tip (1 pixel of the bin4 tomogram = 1.408 nm), and 2-pixel bins for the distance along the tip to minimize the number of empty bins in the 2D histograms (Extended Data Fig. 6a-d).

#### Rim swelling analysis

Based on the 2D histograms of the intermembrane distance, we further analyzed the rim shapes by searching for minimum and maximum peaks with a prominence of at least 1 pixel in the histogram rows, smoothed slightly with a Savitzky-Golay filter (window length 9, polynomial order 2). We defined the peak detection frequency as fraction of complete histogram rows in which a peak was detected, a histogram of peak detection frequencies is shown in Extended Data Fig. 6e. The rim dilation factor was defined as ratio of the mean maximum peak height and the base distance, the intermembrane distance in the back of each rim, calculated by averaging the intermembrane distance values of all points further from the tip than 120 nm, where the rims are not dilated or constricted anymore.

#### Bending energy

The Helfrich bending energy is defined as

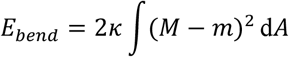

for a membrane with bending rigidity κ, mean curvature M, spontaneous curvature m – assumed to be zero here – and area A^79^. For the discretized n cells of the meshes produced from the segmented rims, we thus calculated the bending energy^80^ from the local mean curvatures M_i_ and cell areas A_i_ as:

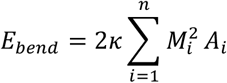

Given that autophagic membranes are known to have high levels of unsaturated lipids^42^, we assumed a bending rigidity κ of 10 k_B_T for the calculations. For a comparison of different rim segments, the resulting energies were normalized by division through the tip length of the rim segment. This normalization should be sufficient since the main contribution to the bending energy comes from the tip area and the contribution in the back is close to zero. To analyze the effect of rim swelling on the bending energy, we generated a reference mesh for each rim mesh with the same overall shape, membrane area and tip length, but with no dilation and a half-toroid-like tip structure. The curvature and bending energy of the reference meshes were determined in the same way as for the original rim meshes.

## Data availability

The data that support this study are available from the corresponding authors upon reasonable request. Source data will be provided upon publication and can be requested by the reviewers if needed.

## Code availability

Any code used in this study is available from the authors upon request.

## Acknowledgement

We thank J. Plitzko for microscope support, L. Bas, B. Engel, W. Wietrzynski, S. Klumpe, S. Khavnekar, R. Bhaskara, S. von Bülow for discussions, and D. Hollenstein and C. Kraft for critical reading of the manuscript. This study was supported by the Max Planck Gesellschaft and funded in part by Aligning Science Across Parkinson’s ASAP-000282 through the Michael J. Fox Foundation for Parkinson’s Research (MJFF). A.B. was supported by a PhD fellowship of the Boehringer Ingelheim Fonds.

## Author contributions

Initial conceptualization, A.B., C.C. and F.W.; methodology, A.B., C.C., P.S.E., F.B., B.A.S., W.B. and F.W.; investigation, A.B., C.C., P.S.E., F.B., F.F., C.W.L., D.L. and F.W.; resources, A.B., C.C., F.F., C.W.L, F.W.; writing – original draft, A.B., C.C., and F.W.; writing – review & editing, A.B., C.C., P.S.E., F.F., C.W.L., D.L., B.A.S., W.B. and F.W.; supervision F.W., P.S.E., B.A.S., and W.B.; funding acquisition, B.A.S and W.B.

## Declaration of interest

B.A.S. holds additional appointments as an Honorary Professor at Technical University of Munich, Germany and adjunct faculty at St. Jude Children’s Research Hospital, Memphis, TN, USA and is on the Scientific Advisory Boards of Interline Therapeutics and BioTheryX. BAS is co-inventor of intellectual property related to DCN1 inhibitors (unrelated to this work) licensed to Cinsano. W. B. holds additional appointments as honorary Professor at the Technical University Munich and Distinguished Professor at ShanghaiTech University and is a member of the Life Science Advisory Board of Thermo Fisher Scientific.

## Notes

### Competing Interest Statement

The authors have declared no competing interest.

