## Supplementary information for "In situ structural analysis reveals membrane shape transitions during autophagosome formation"

1 - Max Planck Institute of Biochemistry, Molecular Structural Biology, 82152 Martinsried, Germany
2 - Max Planck Institute of Biochemistry, Molecular Machines and Signaling, 82152 Martinsried, Germany
3 - Aligning Science Across Parkinson's (ASAP) Collaborative Research Network, Chevy Chase, MD, 20815, United States
4 - Human Technopole, Viale Rita Levi Montalcini, 1, 20157 Milan, Italy
5 - Max Planck Institute of Biophysics, Mechanisms of Cellular Quality Control, 60439 Frankfurt a. M., Germany
6 - Max Planck Institute of Biophysics, Department of Theoretical Biophysics, 60438 Frankfurt a. M., Germany

[#] These authors contributed equally to this work
* Corresponding authors:





#### Extended Data Figures

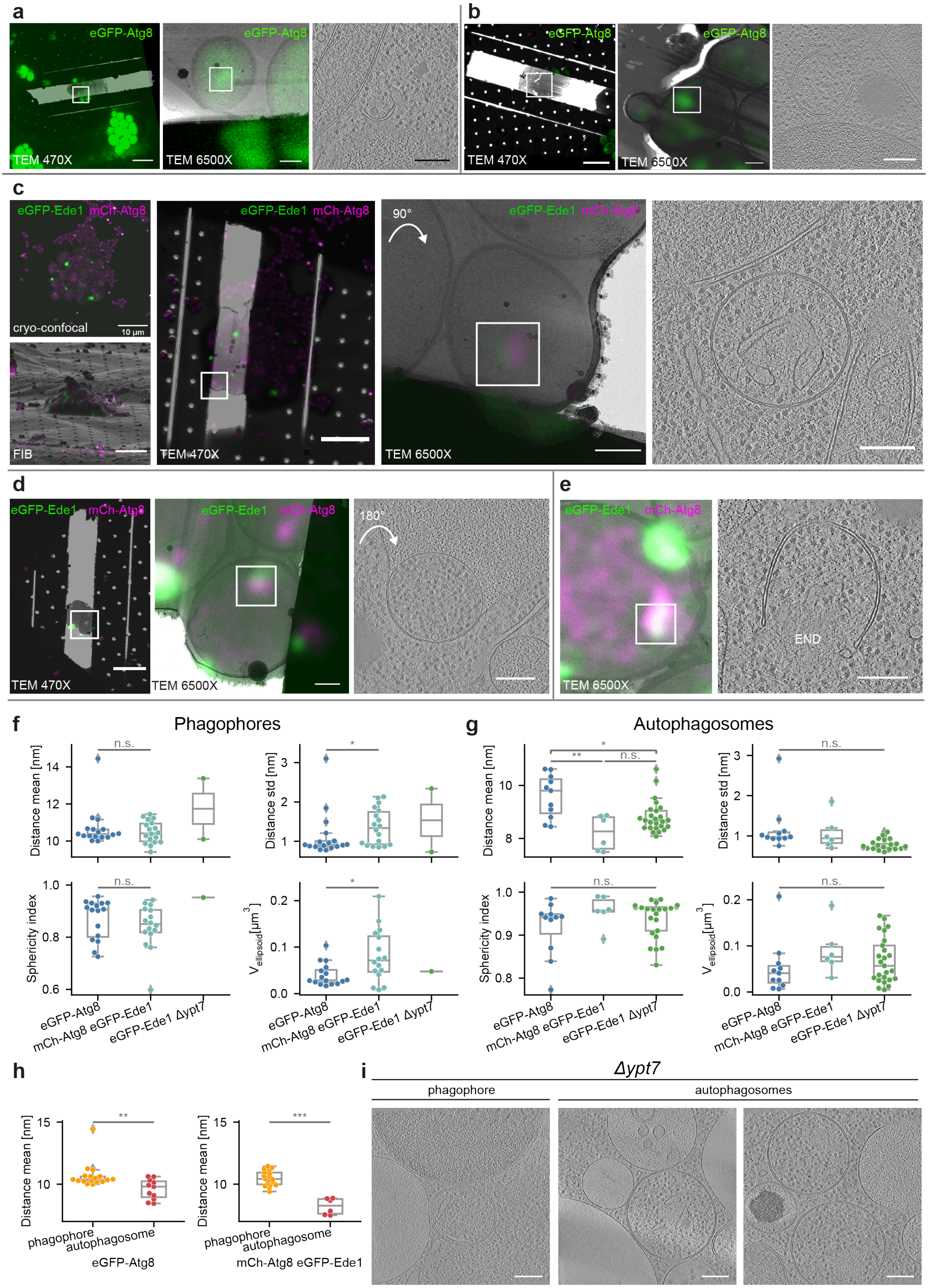

◄**Extended Data Fig. 1. Autophagic structures revealed by correlative cryo-ET. a**, Cryo-fluorescence overlays on TEM maps at 470X and 6500X magnification for the phagophore shown in Fig. 1e. In this case fluorescence stacks were acquired on the final lamella by using a fluorescence microscope integrated in FIB chamber (METEOR). **b**, Cryo-fluorescence overlays on TEM maps for the phagophore in Fig. 1f. **c**, Complete correlative workflow shown exemplary for the autophagosome shown in Fig. 1g. Fiducial-based registration of cryo-fluorescence volume data (projection image in upper left panel) is used to target FIB milling (bottom left) and tomograms acquisition (TEM overviews). **d**, Cryo-fluorescence overlays on TEM maps for fusion structure shown in Fig. 1h. **e**, Example of phagophore captured by using colocalization of eGFP-Ede1 and mCherry-Atg8 signal. **f-g**, Comparison of structures from the three different yeast strains (eGFP-Atg8 (FWY0002), mCherry-Atg8 eGFP-Ede1(FWY0085), and eGFP-Ede1 *Δypt7* (FWY0154)) employed in this study. Plots show mean intermembrane distance, standard deviation of the intermembrane distance, sphericity index and volume of the best fitting ellipsoid for phagophores (f) and autophagosomes (g). **h**, Mean intermembrane distance of phagophores and autophagosomes compared separately for eGFP-Atg8 and EGFP-Ede1 mCherry-Atg8 strains. **i**, Gallery of autophagic structures found in eGFP-Ede1 *Δypt7* strain. All scale bars: 470X, 10 μm; 6500X, 1 μm; tomograms, 200 nm. Statistical analysis: f and h, Mann-Whitney-U test, g: Kruskal-Wallis and pairwise Games-Howell post hoc test. ***$p<0.001$, **$p<0.01$, *$p<0.05$, n.s.: $p\geq0.05$.

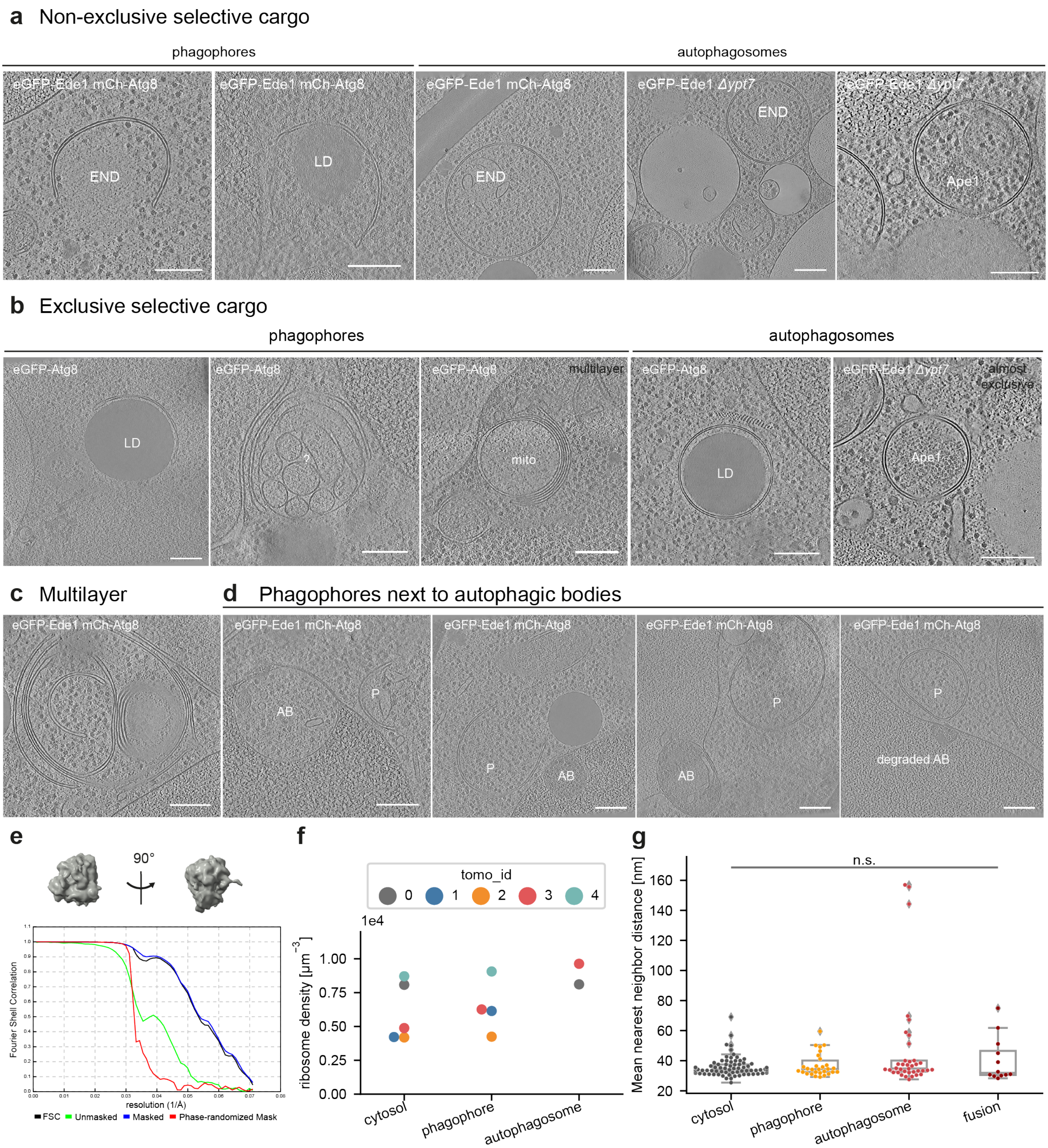

**Extended Data Fig. 2. Autophagic cargo under nitrogen starvation in yeast. a**, Gallery of autophagic structures engulfing selective cargo non-exclusively, i.e., together with cytosol. Last on the right example of structure engulfing Cvt cargo prApe1. All scale bars 200 nm. **b**, Gallery of autophagic structures engulfing exclusive cargo. "?" = unknown membrane cargo. **c**, Example of multilayer structure in which multiple autophagic structures are enwrapping each other. In the center an unperturbed open phagophore engulfing ribosomes. **d**, Examples of open phagophores close to wrapped autophagic bodies (AB) in the vacuole. The last tomogram shows a putative remnant of a degraded AB. **e**, Snapshots and FSC curve of ribosome average (bin2, nominal pixel size 7.04 Å, resolution 15.1 Å) generated from the tomograms. **f**, Ribosome densities (count/volume) determined in 5 example tomograms in the cytosol vs inside phagophores or autophagosomes. **g,** Comparison of mean ribosome nearest neighbor distances in different compartments. Differences between cytosol and autophagic ribosome distances were analyzed with the Wilcoxon signed-rank test, treating values from compartments in the same tomogram as paired (n=77 tomograms).

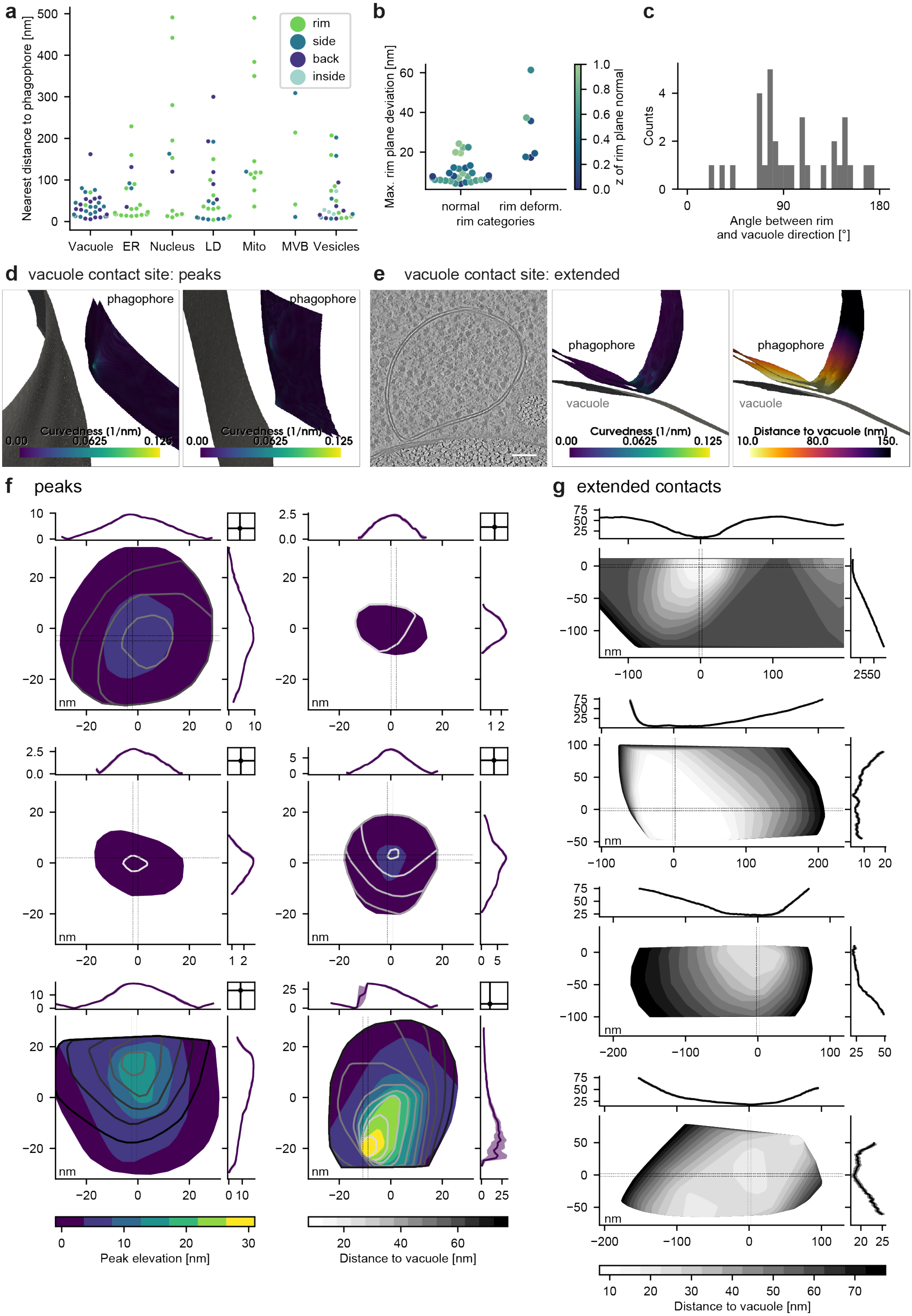
a
Nearest distance to phagophore [nm]
rim
side
back
inside
Vacuole
ER
Nucleus
LD
Mito
MVB
Vesicles
b
Max. rim plane deviation [nm]
normal
rim deform.
rim categories
z of rim plane normal
c
Counts
Angle between rim and vacuole direction [°]
d vacuole contact site: peaks
phagophore
Curvedness (1/nm)
e vacuole contact site: extended
vacuole
Distance to vacuole (nm)
f peaks
g extended contacts
nm
Peak elevation [nm]
Distance to vacuole [nm]

◄**Extended Data Fig. 3. Analysis of phagophore-organelle contact sites. a**, Phagophore interaction area vs distance to different organelles. The color of each point indicates which phagophore area was closest to that single organelle. **b**, Maximum distance of rim points from the best-fitting rim plane for phagophores with an obvious rim deformation vs other phagophores. While larger deviations in some “normal” rims can be explained by difficulties in segmentation caused by the missing wedge if the rim plane is close to parallel to the xy plane of the tomogram, the “rim deformation” examples show large deviations even with the rim clearly visible in all slices. **c**, Histogram of angles indicating the orientation of the rim opening with respect to the vacuole. Angles are calculated between the normal of the phagophore rim plane (pointing outwards) and the shortest phagophore-vacuole vector. **d**, Two examples of peak contact sites between vacuole (left) and phagophore (right). **e**, Example of an extended contact site between an open phagophore (opening not visible in this slice) and the vacuole. **f**, 2D maps and 1D profiles of peak contact sites. Colored peak elevation maps are overlaid with lines indicating the distance to the vacuole. **g**, 2D maps and 1D profiles of extended contact sites. Grey scale maps indicate the distance to the vacuole.

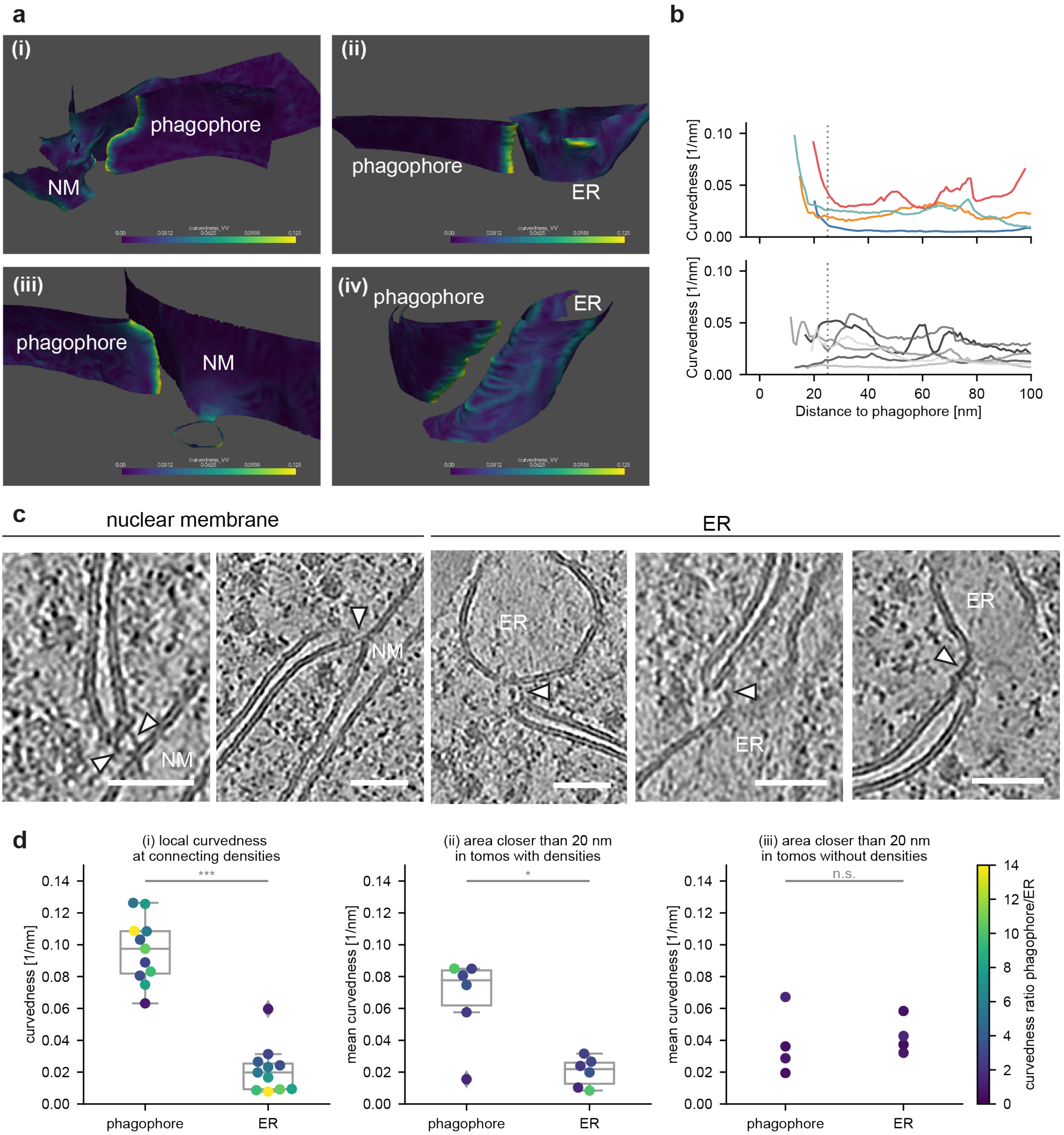

**Extended Data Fig. 4. Analysis of phagophore contact sites with the ER or NM.** **a**, 3D views, colored by curvedness, of nuclear membrane (NM, left) and ER (right) contact sites with phagophore rims. Deformations of the phagophore rim are clearly visible in the NM-phagophore contact sites. **b**, Local ER curvedness vs distance to the phagophore at ER-phagophore contact sites. The upper plot shows examples in which the ER curvedness increases at the contact site, the lower plot shows all other analyzed examples. The dotted line indicates the cutoff distance (corrected for a membrane thickness of 5 nm) which could potentially be spanned by Atg2. **c**, Gallery of rim contact sites with nuclear membrane or ER. White arrowheads indicate connecting densities. Tomograms denoised with cryo-CARE. Scalebars 50 nm. **d**, (i) Curvedness of the phagophore and ER membrane points closest to the segmented connecting densities; (ii) mean curvedness in the area within 20 nm of ER-phagophore distance (membrane surface to membrane surface) in structures for which connecting densities were observed; (iii) same as (ii) but for structures for which no connecting densities were observed. Differences analyzed with Wilcoxon signed-rank test.

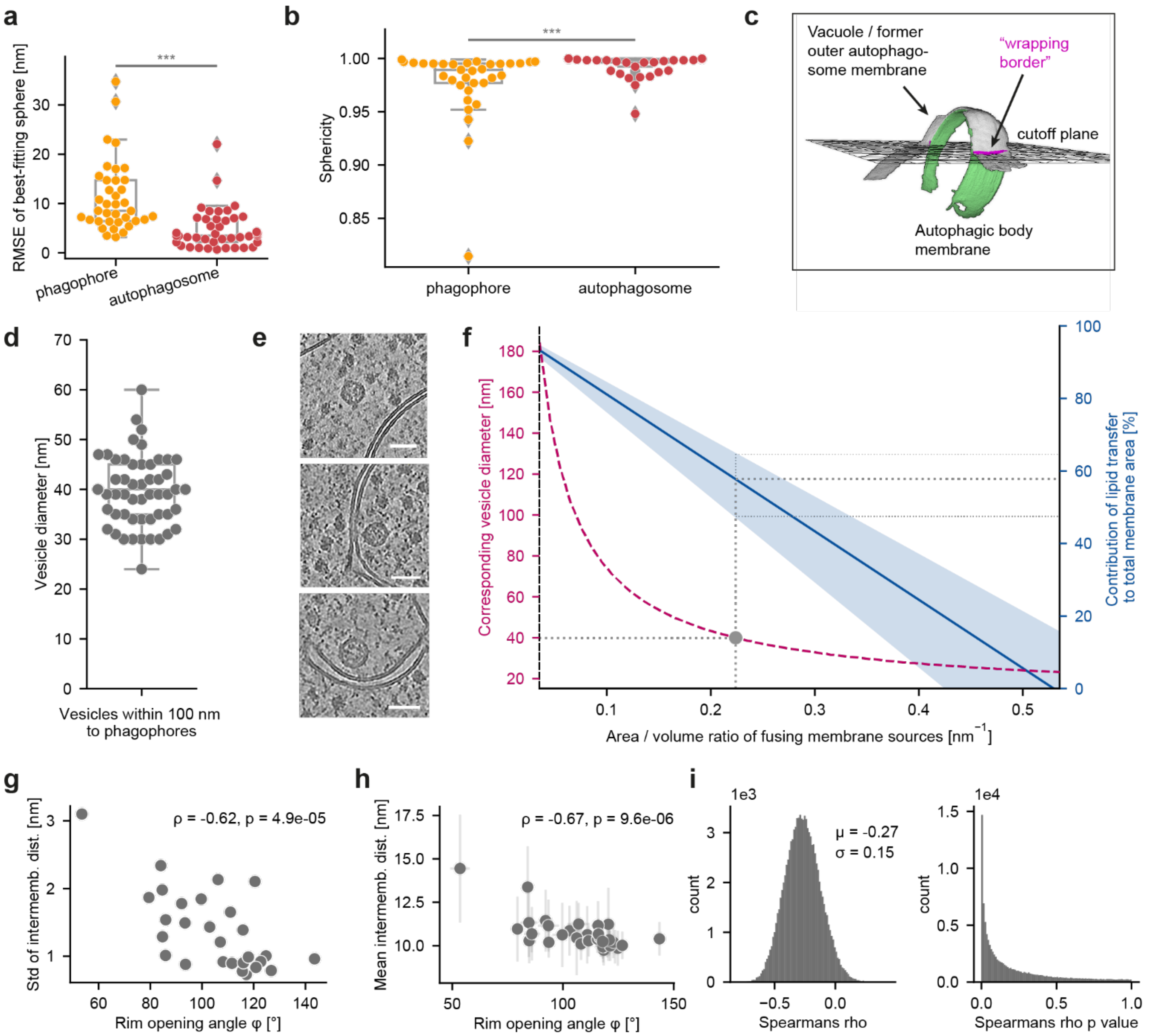

**Extended Data Fig. 5. Analysis of autophagic membrane structures. a-b**, Alternative sphericity measurements confirm that autophagosomes are more spherical than phagophores. **a**, Root Mean Square Error (RMSE) of best fitting spheres to phagophore vs autophagosome inner membranes. **b**, "Classical" sphericity of phagophores and autophagosomes, calculated from best-fitting ellipsoids (fit to inner membrane points) as ratio between the surface area of a sphere with volume equal to the ellipsoid and the surface area of the ellipsoid. **c**, The intermembrane distance of fusion intermediates (Fig. 5c) is measured only for the part wrapped by the vacuole membrane, by cutting the structures with a plane fit through the points at the wrapping border. **d**, Diameters of vesicles observed within 100 nm to the phagophores. **e**, Images of vesicles close to the phagophore membrane. Tomograms denoised with Cryo-CARE. Scalebars 50 nm. **f**, Calculating the contribution of different membrane sources to the autophagosome. Given the combined area/volume (A/V) ratio of all fusing membrane sources (e.g. vesicles) on the x axis, the blue graph indicates the percentage of autophagosome membrane originating from lipid transfer (right y axis), e.g. from the ER. Blue line: calculation with mean autophagosome A/V ratio; blue area: calculation with mean ± standard deviation. For illustration, the magenta line (left y axis) shows the vesicle diameters corresponding to the A/V ratios of fusing membrane sources (x axis). Grey dotted lines indicate calculation results for vesicles with 40 nm diameter. **g**, Standard deviation of intermembrane distance of phagophores plotted over opening angle φ. **h**, Plot showing standard deviations as error bars for the mean intermembrane distance vs opening angle plot showed in Fig. 5f. **i**, Bootstrapping analysis for the correlation between intermembrane distance and φ shown in (**h**): distributions of Spearman's rho and p values obtained by bootstrapping from the raw data.

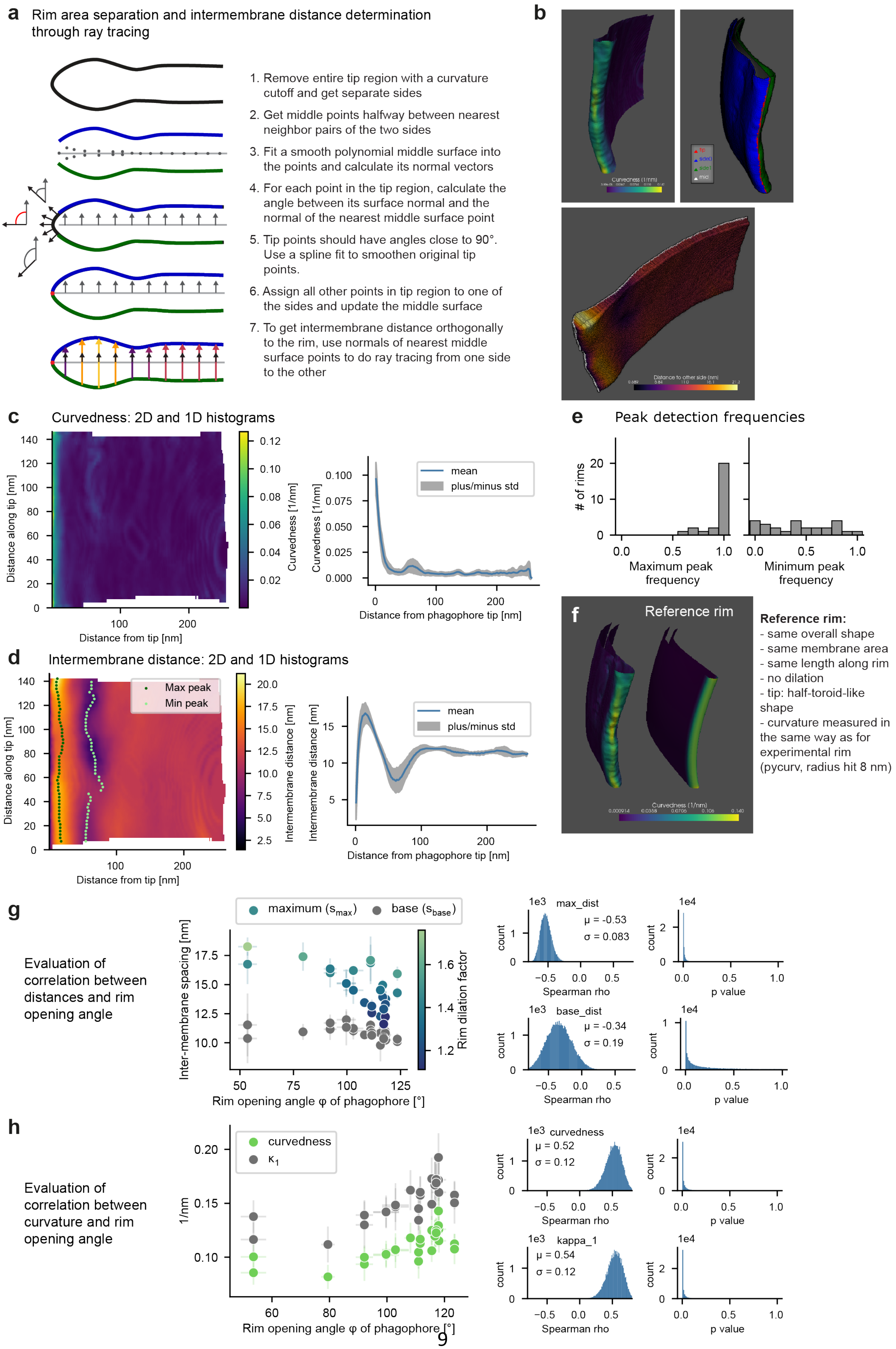
a
Rim area separation and intermembrane distance determination through ray tracing
1. Remove entire tip region with a curvature cutoff and get separate sides
2. Get middle points halfway between nearest neighbor pairs of the two sides
3. Fit a smooth polynomial middle surface into the points and calculate its normal vectors
4. For each point in the tip region, calculate the angle between its surface normal and the normal of the nearest middle surface point
5. Tip points should have angles close to 90°. Use a spline fit to smoothen original tip points.
6. Assign all other points in tip region to one of the sides and update the middle surface
7. To get intermembrane distance orthogonally to the rim, use normals of nearest middle surface points to do ray tracing from one side to the other
b
Curvedness (1/nm)
Distance to other side (nm)
c
Curvedness: 2D and 1D histograms
Distance along tip [nm]
Distance from tip [nm]
Curvedness [1/nm]
mean
plus/minus std
Distance from phagophore tip [nm]
d
Intermembrane distance: 2D and 1D histograms
Max peak
Min peak
Distance along tip [nm]
Distance from tip [nm]
Intermembrane distance [nm]
mean
plus/minus std
Distance from phagophore tip [nm]
e
Peak detection frequencies
### of rims
Maximum peak frequency
Minimum peak frequency
f
Reference rim
Curvedness (1/nm)
Reference rim:
- same overall shape
- same membrane area
- same length along rim
- no dilation
- tip: half-toroid-like shape
- curvature measured in the same way as for experimental rim (pycurv, radius hit 8 nm)
g
Evaluation of correlation between distances and rim opening angle
maximum ($s_{max}$)
base ($s_{base}$)
Inter-membrane spacing [nm]
Rim dilation factor
Rim opening angle φ of phagophore [°]
max_dist
μ = -0.53
σ = 0.083
base_dist
μ = -0.34
σ = 0.19
count
Spearman rho
p value
h
Evaluation of correlation between curvature and rim opening angle
curvedness
$\kappa_1$
1/nm
Rim opening angle φ of phagophore [°]
curvedness
μ = 0.52
σ = 0.12
kappa_1
μ = 0.54
σ = 0.12
count
Spearman rho
p value

◄**Extended Data Fig. 6. Phagophore rim analysis. a**, Steps for automated mapping values on rim and calculation of intermembrane distances by ray tracing orthogonally to the phagophore rim. **c**, Illustration of Rim analysis steps as described in (**a**), shown for same example rim as in (c, d, f). Top left: Mesh of rim segment coloured by curvedness determined with PyCurv (radius hit 8 nm). Top right: Rim separated into tip and two sides, plotted with its corresponding middle surface (white). Bottom: Ray tracing vectors showing local intermembrane distances. **c**, 2D and 1D histograms of the combined curvedness values of inner and outer membrane mapped against the distance from and along the tip for an example rim. **d**, 2D and 1D histograms of mean intermembrane distance vs distance from and along the tip for the example rim. The 2D histogram is used for analysis of frequency and position of maximum and minimum peaks indicating rim dilation and constriction. **e**, Detection frequency of maximum and minimum intermembrane distance peaks in the analyzed rims. A frequency of 1 indicates that a peak was found in every section moving along the tip, while 0.5 indicates that only half of the sections along the tip has a peak. All analyzed rims show maxima (dilation) in at least half of the analyzed segment, whereas minimum peaks (constriction) are not detected consistently. **f**, 3D rendering of a model rim (left) and its corresponding reference rim, coloured by curvedness. Listed on the right are the criteria with which the reference is built. **g**, Maximum and base intermembrane spacing plotted against the rim opening angle φ with error bars indicating standard deviations of mean values. On the right bootstrapping analysis of Spearman correlation coefficients and p values. **h**, Curvedness and first principal curvature $\kappa_1$ at the tip, plotted against φ with error bars indicating standard deviations. On the right, corresponding correlation bootstrapping results.

#### Extended Data Tables

**Extended Data Table 1:** Phagophore peaks at vacuole contact sites, n=7

| | Area [nm$^2$] | Height [nm] | Width [nm] | Min. vac. dist. [nm] | Max. curvedness [nm$^{-1}$] | $\Delta E_{bend}$ [J] | Pearson's ρ (peak elevation vs vac dist) |
|---|---|---|---|---|---|---|---|
| mean | 1916 | 11.5 | 24.2 | 30 | 0.081 | 2.8E-19 | -0.75 |
| std | 1621 | 10.6 | 6.4 | 14 | 0.060 | 4.6E-19 | 0.17 |
| min | 279 | 3.6 | 16.6 | 19 | 0.031 | 1.5E-20 | -0.89 |
| median | 1212 | 7.4 | 23.7 | 23 | 0.061 | 1.0E-19 | -0.80 |
| max | 4088 | 33.2 | 31.9 | 53 | 0.197 | 1.3E-18 | -0.43 |

**Extended Data Table 2:** ER-phagophore contact site analysis, n=10

| | Min. dist. ph-ER [nm] | ER contact area [nm$^2$] | ph contact area [nm$^2$] | ph. mean curvedness [nm$^{-1}$] | ER mean curvedness [nm$^{-1}$] |
|---|---|---|---|---|---|
| mean | 16.4 | 3261 | 3108 | 0.055 | 0.029 |
| std | 3.8 | 4248 | 4027 | 0.028 | 0.015 |
| min | 10.9 | 323 | 265 | 0.016 | 0.008 |
| median | 16.7 | 1609 | 1540 | 0.062 | 0.029 |
| max | 22.7 | 14563 | 13155 | 0.085 | 0.058 |

**Extended Data Table 3:** Rim dilation characteristics, n=26

| | $d_{max}$ mean [nm] | $d_{max}$ SD [nm] | $pos_{max}$ mean [nm] | $pos_{max}$ SD [nm] | $d_{base}$ mean [nm] | $d_{base}$ SD [nm] | Dilation factor |
|---|---|---|---|---|---|---|---|
| mean | 14.7 | 0.78 | 17.0 | 4.3 | 10.9 | 0.72 | 1.35 |
| std | 1.8 | 0.52 | 7.0 | 6.8 | 0.5 | 0.60 | 0.15 |
| min | 11.6 | 0.19 | 10.1 | 0.7 | 9.8 | 0.24 | 1.11 |
| median | 14.5 | 0.61 | 15.6 | 2.6 | 10.8 | 0.47 | 1.32 |
| max | 18.3 | 2.04 | 47.2 | 36.9 | 12.0 | 2.70 | 1.76 |

**Extended Data Table 4:** Curvature at the rim tip, n=26

| | $\kappa_1$ mean [$nm^{-1}$] | $\kappa_1$ SD [$nm^{-1}$] | $\kappa_2$ mean [$nm^{-1}$] | $\kappa_2$ SD [$nm^{-1}$] | Curvedness mean [$nm^{-1}$] | Curvedness SD [$nm^{-1}$] |
|---|---|---|---|---|---|---|
| mean | 0.153 | 0.018 | 0.020 | 0.024 | 0.111 | 0.013 |
| std | 0.019 | 0.003 | 0.014 | 0.008 | 0.014 | 0.002 |
| min | 0.112 | 0.013 | -0.001 | 0.009 | 0.082 | 0.009 |
| median | 0.154 | 0.017 | 0.020 | 0.023 | 0.110 | 0.013 |
| max | 0.193 | 0.023 | 0.060 | 0.037 | 0.143 | 0.018 |

**Supplementary Note 1: Intermembrane distance algorithm**

For an automated analysis of the distance between two more or less parallel surfaces, the two easiest conceivable parameters are the minimum distance between points in the two surfaces (assuming sufficient sampling) and the distances calculated by following the normals of one surface until they intersect with the second surface (normal distance). As described by Kim and colleagues[1], both approaches can fail in the presence of peaks or holes in one membrane. Moreover, calculating consistent normals would necessitate refined segmentations[2]. Based on the algorithm described by Kim et al., we devised a refined minimum distance algorithm that only uses minimum point distances and is robust against peaks and holes in either of the two measured surfaces. The major steps for finding the point-wise distances of a membrane A to a membrane B are (for illustrations see Supplementary Note Fig.1):

1. For each point in A, find the nearest neighbor in B and record the distance (ii).
2. If a point in B was chosen by several points in A, keep the shortest connection and discard all other A-B pairs and their distances (ii).
3. For each point in B, find the nearest neighbor in A.
4. If a point in A that was discarded in step 2 is found as nearest neighbor by a point in B (step 3), add back the B-A pair and its distance to the final set of refined points and distances (iii+iv).

We tested this algorithm on various 2D examples and membrane segmentations and found that it is robust to peaks, holes and overhangs. All distance values reported in Fig. 2 were calculated in this way.

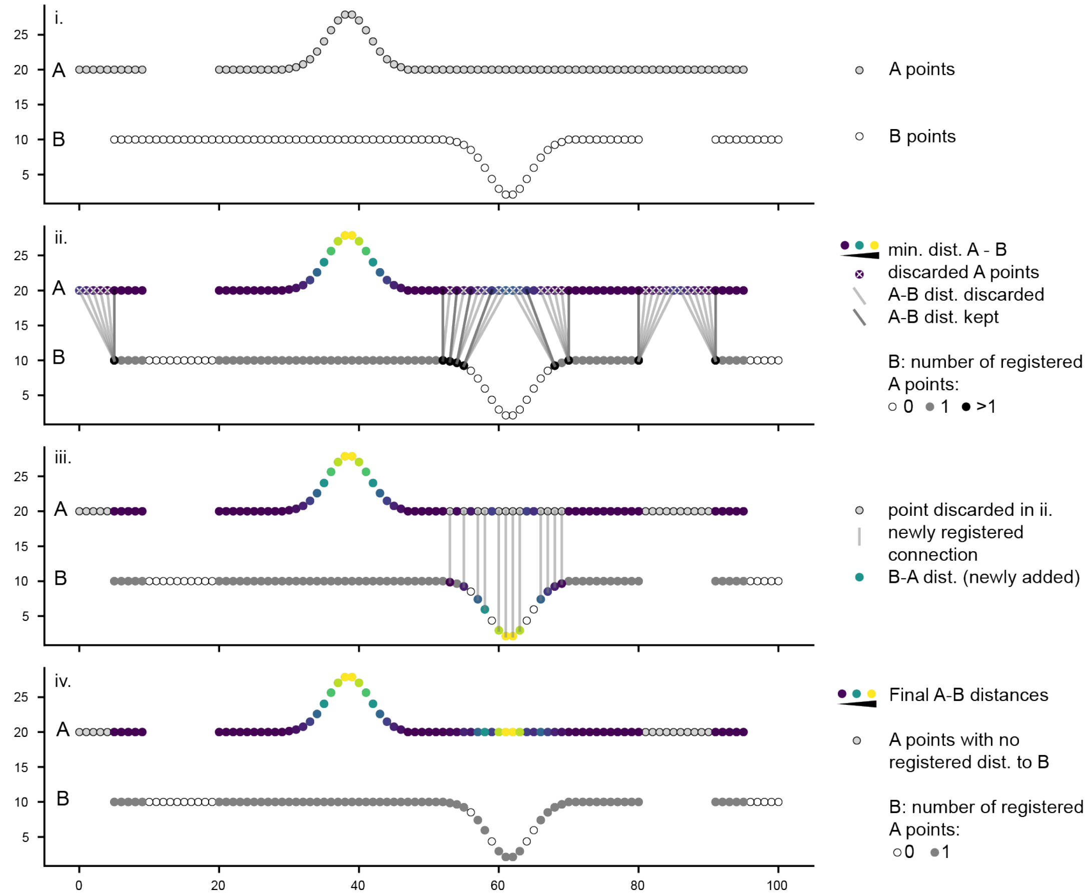

**Supplementary Notes Fig. 1:** Illustration of intermembrane distance algorithm.

**Supplementary Note 2: Estimating the completeness of phagophores.**

One challenge in the analysis of phagophores was to find a parameter to estimate robustly the completeness of each observed structure. First, one could use an ellipsoid fit, cut it with a plane through the rim points and calculate e.g. the fraction of surface areas of the cut vs. complete ellipsoid (Fig. S8a, strategy A). However, especially for early phagophores, ellipsoid fits did not always converge to reasonable final dimensions, even though the used iterative ellipsoid fitting algorithm is more robust to noise than a simple least-squares fit[3]. We thus searched for a parameter that does not rely on an ellipsoid fit, excluding also the bending angle which is used e.g. in Sakai et al.[4] since this would require finding the center of the phagophore which would also necessitate an ellipsoid fit (Supplementary Notes Fig. 2a, strategy B).

In the end, the most robust parameter that we identified is the "rim opening angle" φ, defined as the angle between a plane through the phagophore rim and tangential planes to the phagophore membrane close to the rim. This angle should increase from 0° in an initial membrane disk to 180° just before phagophore closure (Supplementary Notes Fig. 2a, strategy C).

An alternative parameter that does not rely on ellipsoid fits uses circle fits in planes parallel to the rim plane instead (Supplementary Notes Fig. 2a, strategy D). For each phagophore, a plane parallel to the rim plane is moved from the rim towards the back in a stepwise fashion, and at each plane position, a circle is fit to the phagophore points close to the plane. The final parameter reported for each phagophore is the ratio of the maximum radius – the "belly" radius of the phagophore – divided by the radius at the rim. While this ratio should always equal 1 for initial phagophores, as soon as the phagophore reaches its characteristic cup shape, it should increase gradually until closure of the phagophore. After calculating the ratio for all phagophores, we first confirmed that it correlates positively with the rim opening angle (Supplementary Notes Fig. 2b). As in the case of the rim opening angle, the intermembrane distance of phagophores correlates negatively with the ratio of maximum radius and rim radius (Supplementary Notes Fig. 2c). The decrease of intermembrane distance during growth of the phagophore is thus detected through two independent parameters.

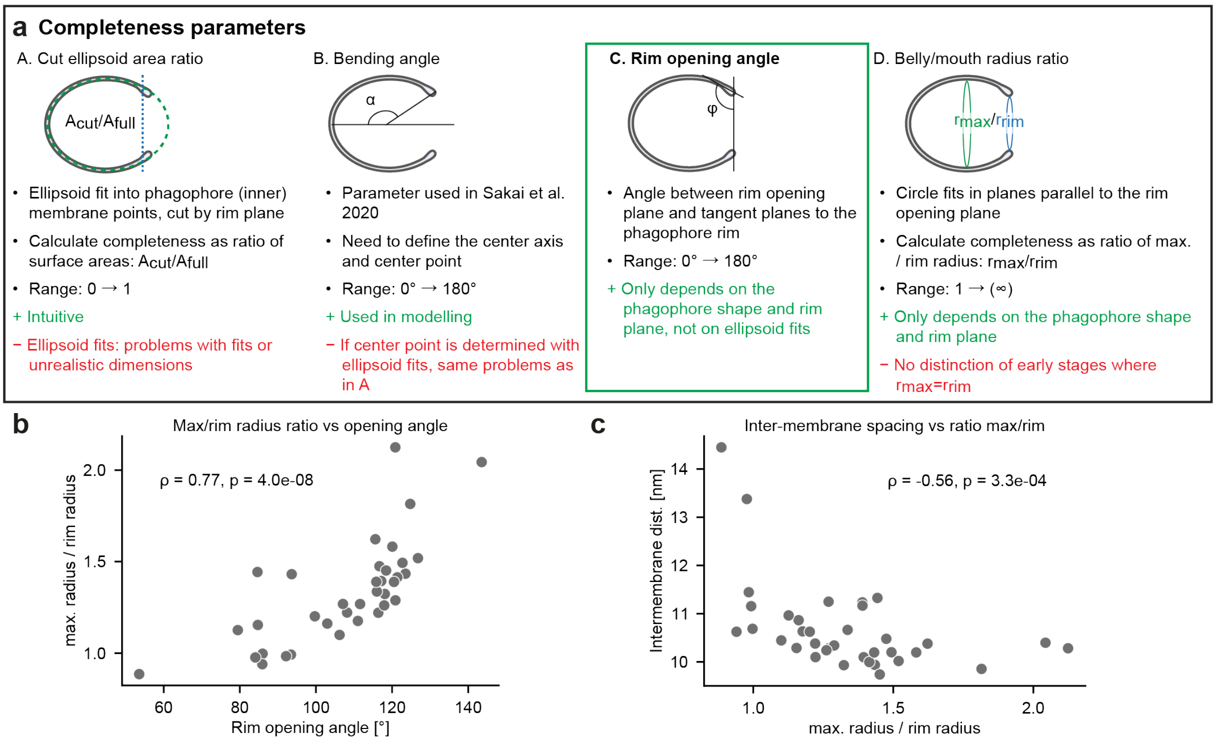

**Supplementary Notes Fig. 2:** **(a)** Comparison of different completeness parameters. **(b)** Correlation of belly/mouth radius ratio (strategy D) with rim opening angle (strategy C). **(c)** The phagophore intermembrane distance correlates negatively with the belly/ mouth radius ratio. Correlation analysis with Spearman's rank correlation.

**Supplementary Note 3: Estimating the number of Atg2 molecules**

To estimate the number of Atg2 molecules needed to build an autophagosome, we follow the argumentation presented in von Bülow & Hummer[5], inserting however the autophagosome dimensions obtained from the tomograms. For an average-sized autophagosome from the experimental data with a lipid bilayer area of 1.6 $\mu m^2$, an area per lipid[6] of 0.724 $nm^2$ and 58% of membrane contribution by lipid transfer (calculation with 40 nm vesicles), around 2.5 million lipids would need to be shuttled through Atg2 into the phagophore. Based on lipid transfer experiments from different groups[7,8] the lipid transfer rate of Atg2 was estimated as 115 lipids/second or 750 lipids/second[5]. Assuming a chemical potential difference between phagophores and ER of 1 $k_BT$ and ten minutes of transfer time[5], the 2.5 million lipids could be transferred by 77 Atg2 molecules for the slow transfer rate, and only 12 Atg2 molecules for the fast transfer rate.

**Supplementary Tables**

**Supplementary Table 1**: yeast strains used in this study

| **Name** | **Relevant genotypes** | **Reference** |
|---|---|---|
| FWY0001 | *MATα, his3-Δ200, leu2-3,2-112, lys2-801, trp1-1(am), ura3-52* | Wilfling et al. 2020[9] |
| FWY0002 | *natNT2::pADH::EGFP::Atg8* | Wilfling et al. 2020[9] |
| FWY0085 | *natNT2::pADH::EGFP::Ede1, pRS305::pADH::mCherry-Atg8* | Wilfling et al. 2020[9] |
| FWY0154 | *natNT2::pADH::EGFP::Ede1, ypt7Δ::hphNT1* | in this study |
| FWY0155 | *pRS305::pADH::mCherry-Atg8* | in this study |
| FWY0156 | *pRS305::pADH::mCherry-Atg8, Vph1::EGFP::kanMX4* | in this study |
| FWY0157 | *pRS305::pADH::mCherry-Atg8, Idh1::EGFP::HIS3MX6* | in this study |
| FWY0158 | *pRS305::pADH::mCherry-Atg8, Sec16::EGFP::HIS3MX6* | in this study |
| FWY0159 | *pRS305::pADH::mCherry-Atg8, Sec61::EGFP::HIS3MX6* | in this study |
